# Re-engineering a lost trait: *IPD3*, a master regulator of arbuscular mycorrhizal symbiosis, affects genes for immunity and metabolism of non-host Arabidopsis when restored long after its evolutionary loss

**DOI:** 10.1101/2023.03.06.531368

**Authors:** Eli D. Hornstein, Melodi Charles, Megan Franklin, Brianne Edwards, Simina Vintila, Manuel Kleiner, Heike Sederoff

## Abstract

Arbuscular mycorrhizal symbiosis (AM) is a beneficial trait originating with the first land plants, which has subsequently been lost by species scattered throughout the radiation of plant diversity to the present day, including the model *Arabidopsis thaliana*. To explore why an apparently beneficial trait would be repeatedly lost, we generated *Arabidopsis* plants expressing a constitutively active form of *Interacting Protein of DMI3*, a key transcription factor that enables AM within the Common Symbiosis Pathway, which was lost from *Arabidopsis* along with the AM host trait. We characterize the transcriptomic effect of expressing *IPD3* in *Arabidopsis* with and without exposure to the AM fungus (AMF) *Rhizophagus irregularis*, and compare these results to the AM model *Lotus japonicus* and its *ipd3* knockout mutant *cyclops-4*. Despite its long history as a non-AM species, restoring *IPD3* in the form of its constitutively active DNA-binding domain to *Arabidopsis* altered expression of specific gene networks. Surprisingly, the effect of expressing *IPD3* in *Arabidopsis* and knocking it out in *Lotus* was strongest in plants not exposed to AMF, which is revealed to be due to changes in *IPD3* genotype causing a transcriptional state which partially mimics AMF exposure in non-inoculated plants. Our results indicate that despite the long interval since loss of AM and *IPD3* in *Arabidopsis*, molecular connections to symbiosis machinery remain in place in this nonAM species, with implications for both basic science and the prospect of engineering this trait for agriculture.

## Introduction

Arbuscular mycorrhizae (AM) are formed during symbiosis between host plants and soil fungi. In AM, the plant provides carbon in the form of lipids and sugar to the fungus, and receives water and nutrients in return (Oldroyd, 2013; Rich et al., 2017). AM can also confer resistance to pathogens and abiotic stress (Ceballos et al., 2013; Aliyu et al., 2019; Begum et al., 2019; Ramírez-Flores et al., 2020). AM are conserved in the majority of plant species from a single origin through the present day, and are thought to have aided in the first colonization of land (Wang et al., 2010; Delaux et al., 2013; Genre et al., 2020). However, multiple independent losses of the AM trait and its genetic machinery have occurred in diverse plant clades (Cosme et al., 2018). Why a trait considered to be beneficial has been so often lost is a puzzle important for not only basic understanding of AM symbiosis’ role in plant resilience, but also for potential crop improvement.

*Arabidopsis thaliana* is one of the ∼7% of plant species that lack the ability to form AM and which have not replaced them with an alternative endosymbiosis (Veiga et al., 2013; Brundrett, 2017; Cosme et al., 2018; Radhakrishnan et al., 2020). Other such non-mycorrhizal (nonAM) plants include multiple economically important species in the Brassicaceae and Amaranthaceae (Brundrett, 2017). Proposed reasons for AM loss include changes in root morphology and lifestyle, altered insect and pathogen resistance, and the carbon cost of supporting the symbiont (Sikes, 2010; Field et al., 2012; Brundrett, 2017; Ma et al., 2018; Poveda et al., 2019). Little experimental validation exists for drivers of AM loss in individual species, let alone an overarching explanation for which species lose AM, why, and when. The proximate genetic causes for mycorrhizal loss, however, are quite clear: a shared subset of specific genes is deleted in all independent nonAM evolutions (Delaux et al., 2014; Radhakrishnan et al., 2020).

Gene losses in nonAM plants prominently include the Common Symbiosis Pathway (CSP) that mediates signal perception and transduction (Cope et al., 2019; Radhakrishnan et al., 2020; Genre et al., 2020). Member genes lost in *Arabidopsis* include the cell-surface receptor *SymRK*, the ion channel *DMI1*, the kinase *DMI3*, and the transcription factor *IPD3* (Delaux et al., 2014; Bravo et al., 2016; Radhakrishnan et al., 2020). These genes mediate a signal transduction pathway leading from AMF signal perception to activation of DMI3 by calcium (Demchenko et al., 2004; Pan et al., 2018; Feng et al., 2019). The released calcium activates DMI3, which then phosphorylates the transcription factor IPD3, enabling its DNA binding activity for regulation of downstream targets (Oldroyd, 2013; Pimprikar and Gutjahr, 2018). Outside the CSP, groups of genes involved in lipid flux from plant to AMF and in vesicle trafficking to the arbuscule are also lost (Radhakrishnan et al., 2020).

*IPD3 (Interacting Protein of DMI3)* is notable among lost genes in nonAM plants. While most such genes belong to families with non-AM-specific homologs, *IPD3* does not, suggesting its function is distinct to symbiosis. Genetic work has also demonstrated the powerful role of this gene in turning CSP signaling into regulation of AM response genes. The *ipd3* knockout phenotype is near-complete elimination of mycorrhization (Watts-Williams and Cavagnaro, 2015). Ectopic expression of IPD3 constitutively activated via phosphomimicking (*IPD3^S50D^*) or truncation to the DNA-binding domain (*IPD3^Min^*), however, results in symbiosis-like gene regulation even in the absence of a microbial signal or upstream CSP genes (Singh et al., 2014; Pimprikar et al., 2016).

Here, we express *IPD3* in *Arabidopsis* based on knowledge of its uniquely powerful function in AM. By restoring expression of this gene that was present in the mycorrhizal ancestors of *Arabidopsis* before loss of AM in the Brassicaceae, we apply a novel means of identifying retained or unrecognized connections between AM genetics and other genetic networks conserved in nonAM plants. Characterization of *IPD3*-expressing *Arabidopsis* targets two specific questions. First, does *IPD3* play additional roles outside of its canonically narrow function in AM establishment? Second, does *IPD3* retain functional molecular relationships when restored to *Arabidopsis*, which lost its ancestral copy of this gene along with the trait approximately 65 million years ago (Hohmann et al., 2015)? If so, could reconnecting such relationships alter the response to AMF or even restore symbiosis?

We use transcriptomics to understand the gene-regulatory impact of restoring *IPD3* expression to *Arabidopsis*, and we also compare the inverse case of *IPD3* knockout in a mycorrhizal host plant, via the *cyclops-4* mutant of *Lotus japonicus.* As shown in figure 1, our experiment generates a factorial set of transcriptome data for every combination of *IPD3* genotype, [non]symbiotic species background, and AMF exposure. While *Lotus* has a functioning upstream CSP that normally activates IPD3 in the presence of AMF, leading to AM, the *cyclops-4* mutant lacks functioning *IPD3* and results in *Lotus* that fails to form mycorrhizae despite having the upstream CSP elements intact (Yano et al., 2008; Singh et al., 2014; Pimprikar et al., 2016). This *Lotus* mutant phenotypically mirrors wild type *Arabidopsis* in lacking AM, though *Arabidopsis* further lacks many other genes as described above (Delaux et al., 2013). We additionally examined a set of AMF-exposed *Arabidopsis* plants under low- macronutrient conditions (N, P, K) as a known regulator of symbiotic interactions in host plants (Figure 1C) (Bonneau et al., 2013; Nouri et al., 2014; Carbonnel and Gutjahr, 2014). We perform correlation network and differential expression analyses to identify patterns in transcriptomic data from this experiment, and explore their functional implications using gene ontology (GO).

**Figure 1:**
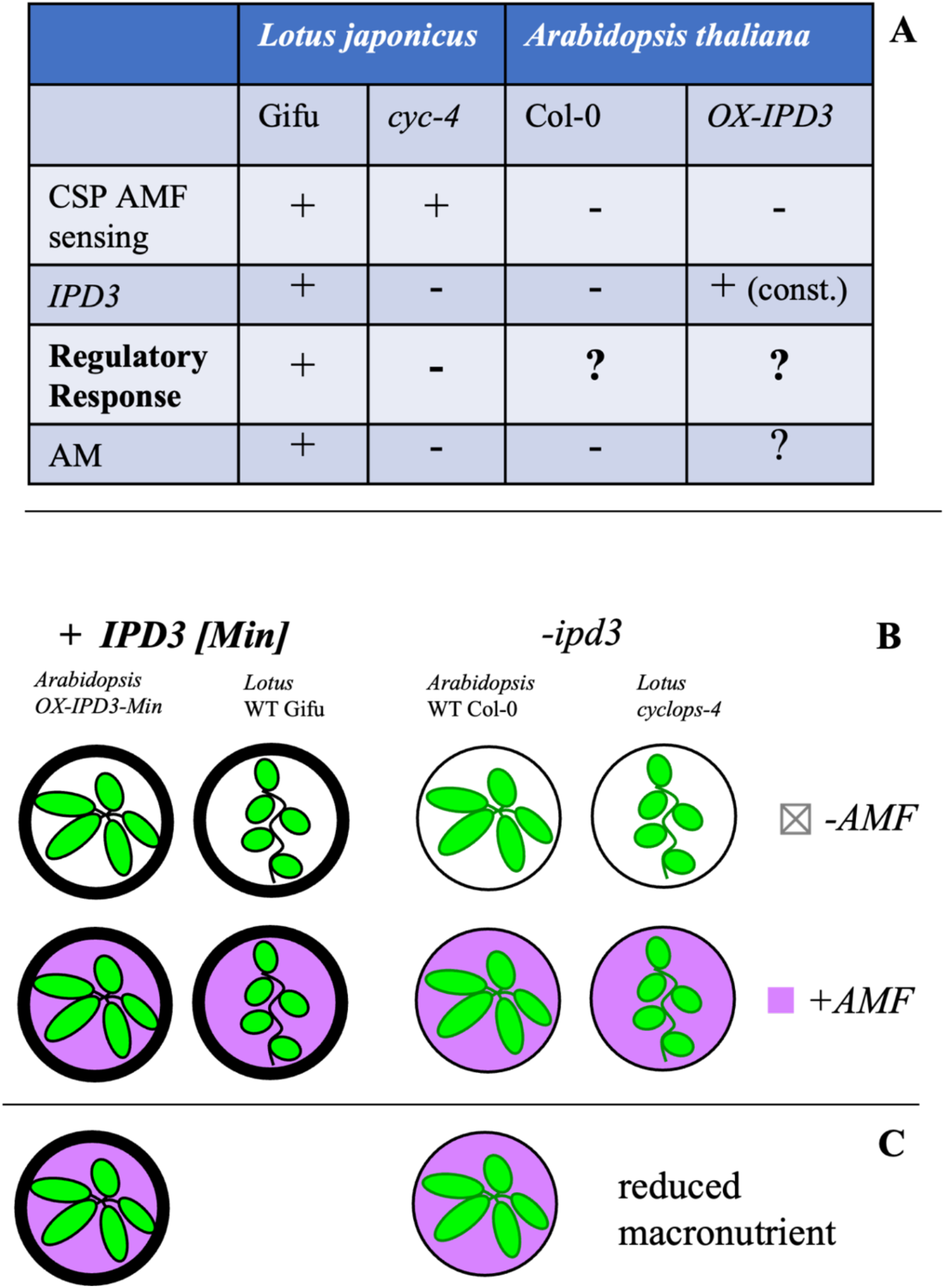
Experimental design comparing different species and genotypes. (A) The AM-host *Lotus japonicus* wildtype (ecotype Gifu) and *IPD3* knockout (*cyclops-4; cyc4*) genotypes differ in their ability to establish AM. *Arabidopsis thaliana* as a non-host species was engineered to express the DNA binding domain of IPD3 (*OX-IPD3^Min^*). (B) Phenotypic and transcriptomic analysis was carried out on the genotypes with or without AM inoculation and (C) under high and low macronutrient-containing media. Two independent transgenic lines of *Arabidopsis* were used for analysis in (B) and 1 transgenic line of *Arabidopsis* was used for analysis in (C).

## Results

### Generation of transgenic *Arabidopsis*

We generated 7 homozygous transgenic *Arabidopsis* lines expressing *Medicago truncatula IPD3* (*IPD3^Mt^*), 4 lines of phosphomimic *IPD3 (IPD3^S50D^*), and 5 lines expressing *IPD3^Min^* (aa 254-513 of IPD3^Mt^) (Singh et al., 2014). All transgenes were expressed under the *Arabidopsis UBIQUITIN10* promoter (Grefen et al., 2010; Ivanov and Harrison, 2014). RT-PCR confirmed expression of all versions of the transgene for 3 generations. Western blot of root and shoot tissue of T3 individuals confirmed presence of IPD3^Min^ protein, but not the full-length versions IPD3^Mt^ and IPD3^S50D^ (Figure S1). We confirmed the identity of extracted IPD3^Min^ using shotgun proteomics of SDS-PAGE gel bands at the expected size (Data S1). *IPD3^Mt^* and *IPD3^S50D^* lines were analyzed in the same manner, however, protein expression was again not detected. We then performed untargeted shotgun proteomics of shoots and roots (n=5). IPD3^Mt^ and IPD3^S50D^ were again not detected, while IPD3^Min^ was highly abundant (top 10% NSAF) in respective lines (Data S1). Consequently, we focused only on characterizing the 5 *IPD3^Min^*lines (numbered 303; 308; 310; 312; 357).

### Phenotypic effect of *IPD3^Min^* expression in *Arabidopsis*

Plants were surveyed for differences in growth under long-day conditions (Figure 2, Figure S2). All 5 transgenic *IPD3^Min^*lines were significantly slower to initiate the transition to flowering than wild type Col-0, measured as days to onset of bolting (Figures 2A&B, Data S2). Four of 5 lines (excluding 303) were also significantly slower to flower than a null segregant control (Figure 2B). Despite flowering differences, no significant difference in seed yield was detected over 3 repetitions of the experiment (Figure 2C). No difference in germination timing was observed (Data S2). Most *IPD3^Min^*transgenic plant roots were bright pink, while color in *IPD3^Mt^* and *IPD3^S50D^* lines was absent or reduced (Figure S3). Nine independent T1 individuals transformed with an empty vector had a range of pink coloration in roots and anthocyanin and carotenoid extraction yielded no measurable amounts (Figure S3). We concluded that the color visible in *IPD3^Min^*roots is the mCherry marker, and variation in intensity is consistent with the other evidence for silencing of the two full-length constructs (Figure S1, Data S1).

**Figure 2:**
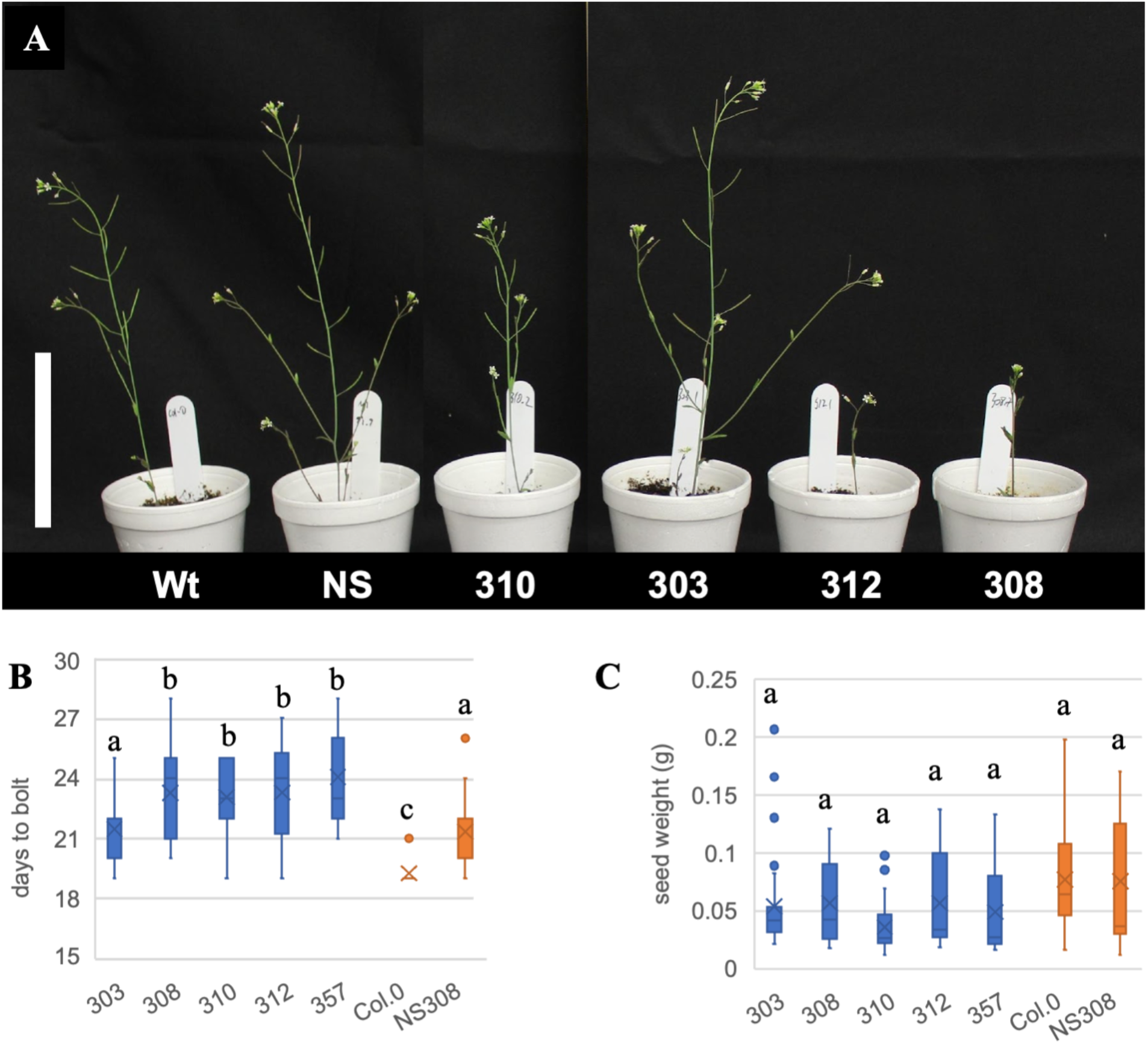
Phenotypes of independent homozygous transgenic lines expressing the DNA-binding domain of IPD3 (IPD3^Min^). (A) Shoots of most *IPD3^Min^* transgenic lines appear shorter, less branched and delayed in development; scale bar 10 cm. (B) *IPD3^Min^*Transgenic lines (blue) had a delayed transition to reproductive development (onset of bolting) relative to nontransgenic controls (orange) (ANOVA, p<0.05, n=15), but (C) showed no significant differences in total seed yield (MCM p<0.05) (in B&C mean is marked with X and median marked with –, outliers shown as dots) (See Data S2).

We also monitored *Lotus japonicus* plants used for cross-species comparison in this experiment for overt differences between the *cyclops-4* knockout mutant and wild type. Consistent with prior literature we noted no obvious growth phenotype in the IPD3 mutant relative to the wild type in *Lotus* (Yano et al., 2008; Horváth et al., 2011) (Figure S4). The genotype of *cyclops-4 Lotus* was confirmed by Sanger sequencing of the *CYCLOPS* genomic region.

### Transcriptional consequences of manipulating *IPD3*

To determine gene regulatory effects of expressing *IPD3^Min^* in *Arabidopsis* and loss of *IPD3* activity in *Lotus*, we sequenced total mRNA of 6-week-old roots of transgenic *IPD3^Min^*, mutant *cyclops-4*, and wild type *Arabidopsis* Col-0 & *L. japonicus* Gifu plants with or without 48 hours of AMF germinated spore treatment prior to collection (Figure 1). Two independent transgenic lines of *Arabidopsis* (308 & 310, Figure 2) were included in the experiment to control for insertion site effects. Transcriptome analyses of *Arabidopsis* incorporated both transgenic lines. Correlation network construction used quantitative expression of *IPD3^Min^*in each sample as the primary transgenic trait. In differential expression analysis an *IPD3^Min^* category including only genes independently attested as DEGs in both transgenic lines was used. We also compared Col-0 *Arabidopsis* to a single *IPD3^Min^*transgenic line (308) with AMF inoculation on low-nutrient (LN) MS medium containing 0.5% P, 1% N, and 1% K relative to ½ MS used in the primary experiment (Figure 1).

In principal component analysis (PCA) of *Lotus*, Gifu transcriptomes clustered according to AMF treatment along PC1 (Figure S5). *cyclops-4* transcriptomes were separated from those of Gifu plants along PC2. Interestingly, both AMF-treated and untreated *cyclops-4* transcriptomes clustered along PC1 with those of AMF-<u>treated</u> Gifu plants. In *Arabidopsis*, *IPD3^Min^*transgenic lines clustered relative to Col-0 along PC1 (Figure S5). Line 308 clustered farther from Col-0 than line 310, corresponding to 4-5X higher *IPD3^Min^* expression in individuals of this line (Figure S5, Figure S6). Col-0 plants clustered by AMF treatment along PC2, but there was no clear separation by treatment among *IPD3^Min^*lines (Figure S5). PCA of the low-nutrient experiment also shows strong clustering by genotype (Figure S7).

### Correlation network and gene ontology analysis

We used weighted gene co-expression network analysis (WGCNA) as a transcriptome analysis tool which captures global expression patterns reflective of the underlying reality of coordinated expression of many genes affected by multiple factors (Langfelder and Horvath, 2008; Langfelder and Horvath 2012). WGCNA clusters the transcriptome into modules with similar expression profiles which suggest they may regulate each other or are co-regulated. WGCNA also correlates expression of a representative eigengene for each module to traits and treatments, in this case quantitative *IPD3^Min^*expression, AMF treatment, and nutrient level. The two-step process of module construction and module-trait correlation makes WGCNA a more sensitive means of linking expression changes that may be subtle at the individual gene level to experimental variables; this can be used to guide more targeted analysis. Network analysis of *Arabidopsis* identified 22 co-regulatory modules. (Figure S8 and Data S3). Figure 3A includes all modules significantly correlated to *IPD3^Min^*expression, and selected modules correlated to AMF treatment and nutrient level. We used gene ontology (GO) enrichment to link module membership and experimental variables to biological functions (Figure 3B). We also performed targeted pathway analysis of specific gene members within modules to better connect these broad patterns to physiological functions (Figure 4). Statistical significance in Figure 4 reflects a hypothesis test of whether mean expression of the given gene differs between specific treatment groups, an independently calculated and complementary measure to the network analysis correlations which establish module membership.

**Figure 3:**
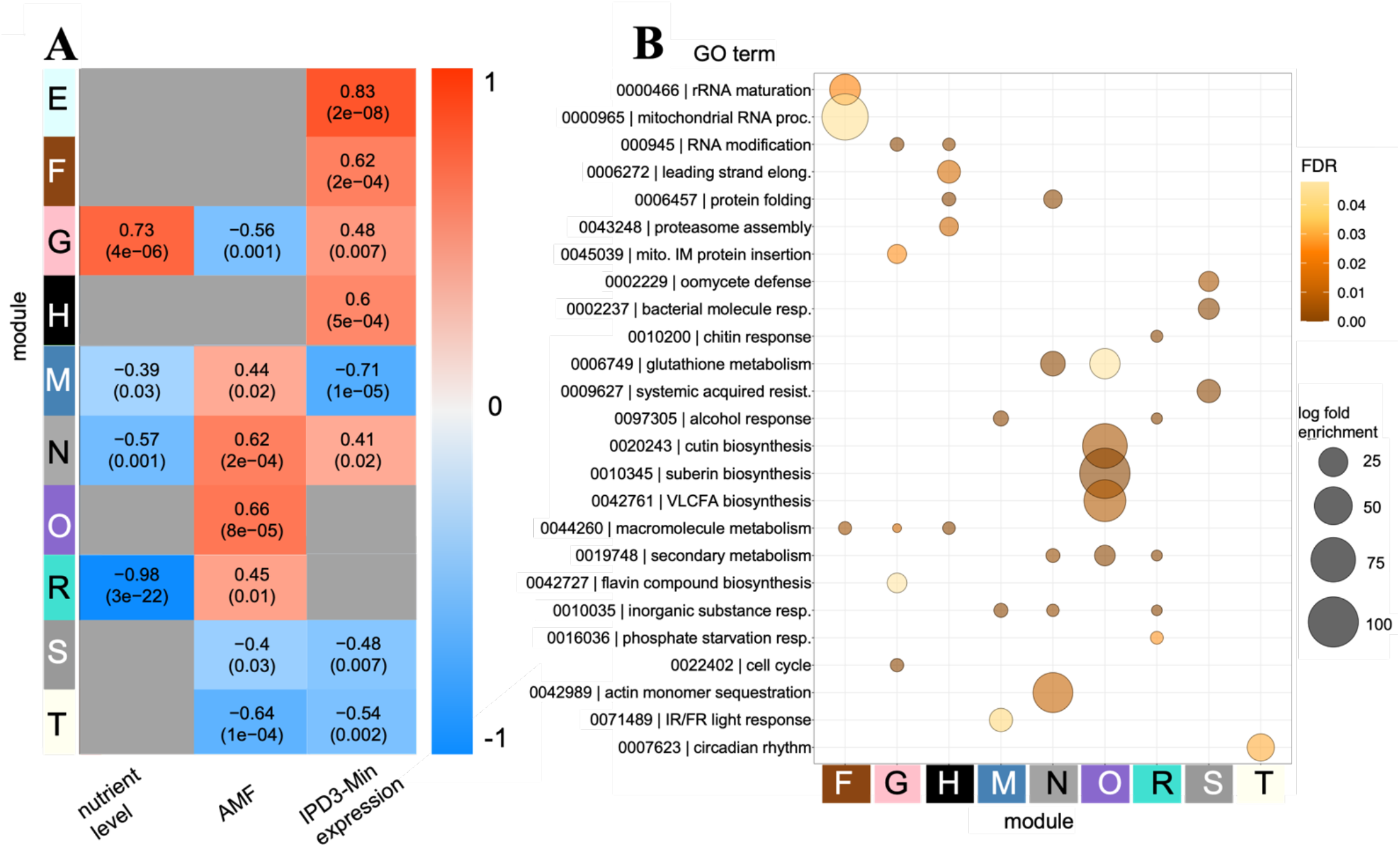
Correlation network modules and gene ontology analysis for transcription of *Arabidopsis thaliana IPD3^Min^* and Col-0 genotypes across AMF and low-nutrient treatments. (A) Selected correlation modules, including all those significantly correlated to *IPD3^Min^*expression. Modules are labeled E-T in order of appearance in the full network (Figure S4). Color of cells within the heatmap reflects correlation of that module to trait values; p-value of the module-trait correlation is listed in parentheses. (B) Gene Ontology enrichment of all modules shown in (A); terms have been reduced by applying a cutoff for the top 3 most-enriched and most-significant terms in each module followed by overlap analysis to establish representative terms. Color intensity corresponds to FDR-corrected p-value of term enrichment and circle area corresponds to scale of enrichment.

**Figure 4:**
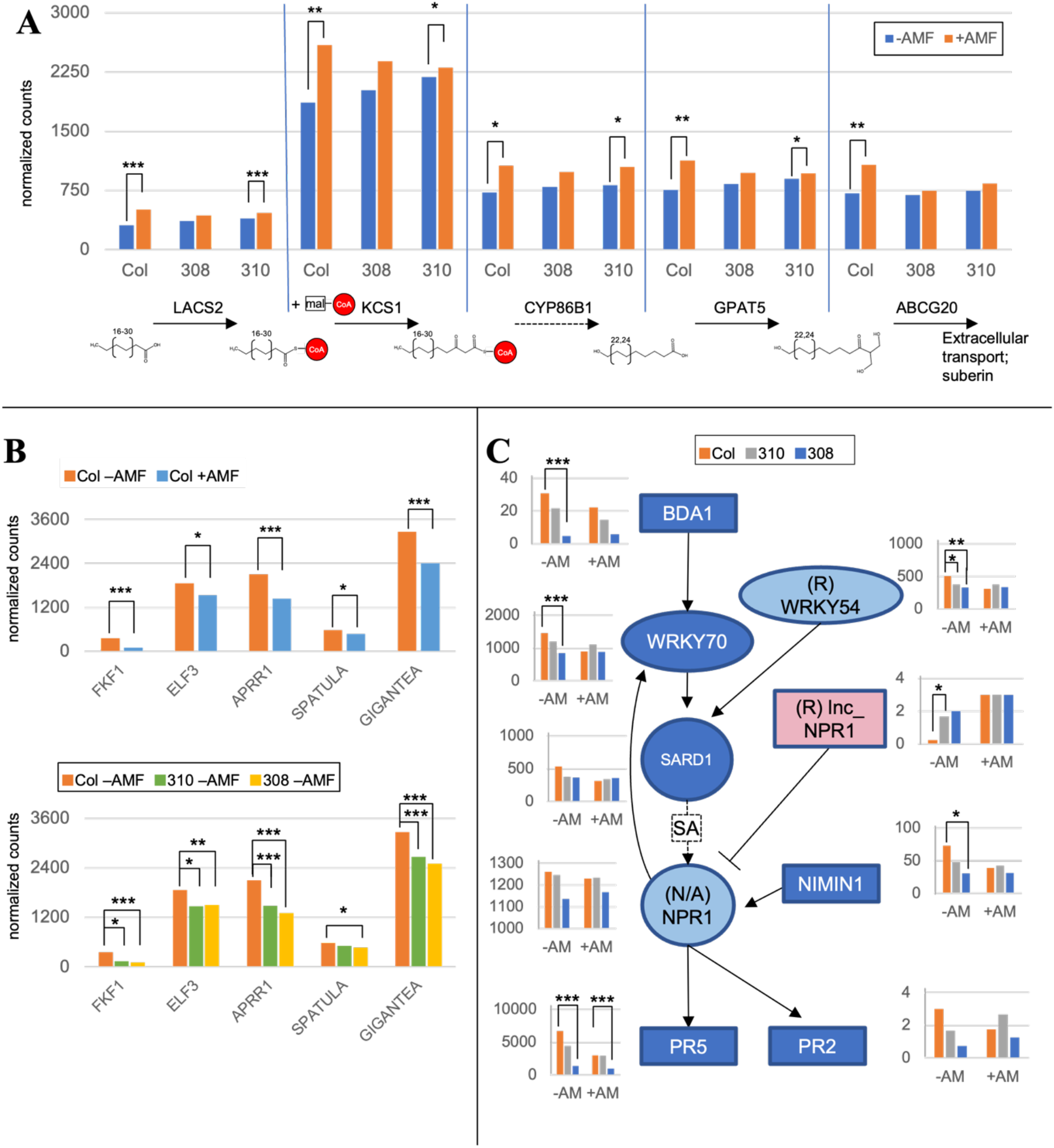
Targeted analysis for pathways of interest within correlation network modules. All genes included in Figure 4 are statistically significant members of the respective network module (having module-trait correlations at p_GS <0.05 as indicated in Figure 3) at p_kME <0.05. Significance reflects 1-tailed t-test of group means for transcript counts of specific genes, a complementary measure which is calculated independently from both the correlation network and untargeted DEG analysis; n=3-5, * = p <0.05; ** = p <0.01; *** = p <0.005. Some interactions in pathway maps have been omitted for clarity. (A) Expression of a subset of individual genes belonging to module O related to extracellular wax synthesis. (B) Expression of well-known circadian clock related genes in module T. (C) Pathway map of the WRKY70-SA-NPR1-mediated defense response with expression of individual genes. Genes belong to module S unless marked otherwise in parentheses

### The *Arabidopsis* response to AMF includes altered defense and lipid metabolism

Despite its nonAM status, *Arabidopsis* transcriptome networks showed 10 modules of co-regulated genes with trait correlations to AMF treatment, indicating a clear perception of and response to the fungus (Figure S4). We considered modules O and R that correlate to AMF treatment but not *IPD3^Min^*expression to reflect portions of the native *Arabidopsis* AMF response that are unaffected by *IPD3^Min^* expression. Expression of genes in module R positively correlated with AMF treatment but negatively correlated with nutrient level. Top GO terms in this module related to both nutrient stress (GO:0016036; phosphate starvation) and biotic interactions (GO:0010200; chitin response) (Figures 4B & S10, Data S4). Given that AMF was the only microbe tested, the biotic interaction terms and linkage to phosphate starvation may represent a generalized microbial response consistent with known effects in *Arabidopsis* (Finkel et al., 2019).

Module O expression correlated solely to AMF treatment, and was highly enriched for terms related to biosynthesis of extracellular lipids (GO:0010143; cutin biosynthethis, GO:0010345; suberin biosynthesis, GO:0042761; very long-chain fatty acid biosynthesis) (Figure 3B). These genes fall along the biosynthetic pathway of 2-monoacylglycerols (2-MAGs), which affect pathogen and stress resistance as components of cutin and suberin (Figure 4A, Data S3, S4). 2- MAGs are also the primary form of carbon exported to AMF fungus from AM host plants (Luginbuehl et al., 2017; Rich et al., 2017). Mean expression of all tested genes in the pathway was significantly higher in AMF-treated Col-0 plants than in untreated plants (Figure 4A).

Significantly higher expression was also detected in some but not all AMF-treated transgenic lines for the same genes. This indicates that upregulation of lipid biosynthetic genes in AMF-treated *Arabidopsis* is *IPD3^Min^-*independent, consistent with the module’s correlation to AMF treatment but not *IPD3^Min^* expression in Figure 3A. Other genes in module O also function in lipid synthesis, including *GPAT6*, a functional homolog of *RAM2* which is strictly required for AM in host species (Data S3) (Gobbato et al., 2013; Dai et al., 2022). While it is a member of module O, increased *GPAT6* expression was only statistically significant for AMF treatment in *IPD3^Min^* line 308, not Col-0 or *IPD3^Min^* 310 (Data S5).

### *IPD3^Min^* expression imitates effects of AMF treatment in non-AMF-treated plants

Transcription in module T correlated negatively to AMF treatment and *IPD3^Min^* expression (Figure 3A). This is consistent with *IPD3^Min^*expression enhancing the effect of AMF treatment or replicating the effect of AMF in untreated samples. A similar pattern was present in modules N (both positive) and S (both negative) (Figure 3A). Module T contains only 56 genes and was enriched solely for terms related to circadian rhythm (GO:0007623; circadian rhythm, GO:0048511; rhythmic process) (Figure 3B, Data S4). Members of this module include components of the *CONSTANS-FT* daylength sensing system (Data S3, S5) (Takagi et al., 2023). As shown in Figure 4B, AMF treatment of Col-0 plants results in significantly reduced expression for genes across the pathway, and *IPD3^Min^* expression replicates this effect even in the absence of AMF.

Module S transcription was negatively correlated with both *IPD3^Min^* and AMF treatment, and is enriched for GO terms related to pathogen defense and systemic acquired resistance (SAR) (GO:0002229, oomycete defense; GO:0002237, bacterial molecule response; GO:0009627, systemic acquired resistance) (Figure 3). This module contains the key defense regulator *WRKY70*, which mediates selection of defense responses by positively regulating salicylic acid (SA) immunity and negatively regulating jasmonic acid (JA) related pathways (Li et al., 2006). As shown in Figure 4C, genes acting up- and downstream of *WRKY70* in SA-mediated defense are also present in this module, including cell-surface ankyrin protein *BDA1*, and pathogen response gene *PR2* (Thomma et al., 2001; Yang et al., 2012a). *NPR1*, another key positive regulator of SA-mediated defense was detected in the transcriptome but not placed in a network module (Figure 4C)(Thomma et al., 2001).

Several SA defense genes (*BDA1; WRKY70; WRKY54; NIMIN1; PR5*) were significantly downregulated in one or both transgenic lines even in the absence of biotic interaction with AMF (Figure 4C). This suggests that *IPD3^Min^* expression in isolation can replicate the suppressive effect of AMF interaction on SA-related defense. Consistent with downregulation of SA-related genes, we also found evidence of antagonistic crosstalk between jasmonic and salicylic acid-mediated defenses (Li et al., 2006; Hou and Tsuda, 2022). Module R, which in contrast to Module S was upregulated in response to AMF, is functionally enriched for jasmonic acid (JA) signaling (GO:009753) (Data S4). JA genes upregulated by AMF in module R included upstream (jasmonate methlytransferase *JMT*) and downstream (chitinases *PR3* and *PR4*) members of the JA defense pathway (Samac et al., 1990; Thomma et al., 1998; Seo et al., 2001). Module R also contained a set of genes enriched for salicylic acid signaling, some of which are negative regulators of SA defense such as At1g08667.1 (*lnc_NPR1*), a putative antisense transcript of *NPR1* whose upregulation would correspond to downregulation of *NPR1* and the SA response (Figure 4C, Data S5). Other SA-related genes were placed into module R due to expression correlation at the individual sample level, but upon comparison of treatment group means matched the pattern of downregulation in module S; this included *WRKY54*, a co-regulator with *WRKY70* of SA synthesis (Chen et al., 2021b) (Figure 4C, Data S5).

Positive *IPD3^Min^* correlation with modules F, G, H, and N corresponded to GO enrichment for background processes, e.g. GO:0000466, rRNA maturation (Figure 3). These functions are essential to the organism, but limited in GO analysis from being linked to specific effects. These modules contained 4,132 genes, indicating that about 10% of all gene models in the *Arabidopsis* genome were upregulated in connection to *IPD3^Min^* (Data S3). Module E had the strongest correlation to *IPD3^Min^*expression but resulted in no significant GO term enrichment (Figure 3A, Data S3, S4). Notably, 8 of 25 genes subsequently identified as *IPD3-*responsive via differential transcription analysis in Figure 6B belonged to module E and are discussed in later sections (Data S6).

### Network analysis of *Lotus* transcriptomes

We also constructed a correlation network for *Lotus*, which is available in Figure S9 and Data S7. A distinctive feature of the *Lotus* network is that while expression of 8 modules correlated with *IPD3* genotypes and 15 modules correlated with the AMF treatment, no modules correlated with both treatments (Figure S9). We confirmed transcription of CSP and AMF marker genes including *CCaMK, NSP2,* and *PT4,* however, all detectable CSP genes were placed into network modules not correlated to either treatment (Data S7).

### Cross-species comparison of differential transcription

Next, we conducted differential transcript abundance analysis for genotype and AMF treatment in *Lotus* and *Arabidopsis*. The intent of this analysis was to characterize transcript abundance changes related to the presence (*Lotus* Gifu, *Arabidopsis IPD3^Min^*) or absence (*Lotus cyclops-4*, *Arabidopsis* Col-0) of the respective *IPD3* versions in a manner that enables cross-comparison of species as directly as possible. The “-*ipd3”* genotypes (Col-0 and *cyclops-4*) and -AMF condition were treated as controls in both species, resulting in an equivalent set of genotype-by-AMF contrasts (Figure 5A). In the *-ipd3 cyclops-4* genotype of *Lotus*, AMF exposure induced only 15 DEGs (Figure 5A contrast A), consistent with prior knowledge that AM symbiosis is strongly *IPD3-*dependent (Yano et al., 2008). In -*ipd3* Col-0 *Arabidopsis*, however, AMF induced 497 DEGs (Figure 5A contrast C). Conversely, AMF treatment of *+IPD3 Lotus* induced 461 DEGs (contrast B) while AMF treatment of *+IPD3^Min^ Arabidopsis* induced only 3 (contrast D). In -AMF *Lotus* the presence of *IPD3* in *+IPD3 Gifu* produced 338 DEGs relative to *-ipd3 cyclops-4* (Figure 5A contrast E), while the equivalent contrast in *Arabidopsis* produced 25 DEGs (contrast F). Contrast of the two genotypes under AMF treatment resulted in similar DEG numbers in both species (contrasts G and H).

**Figure 5:**
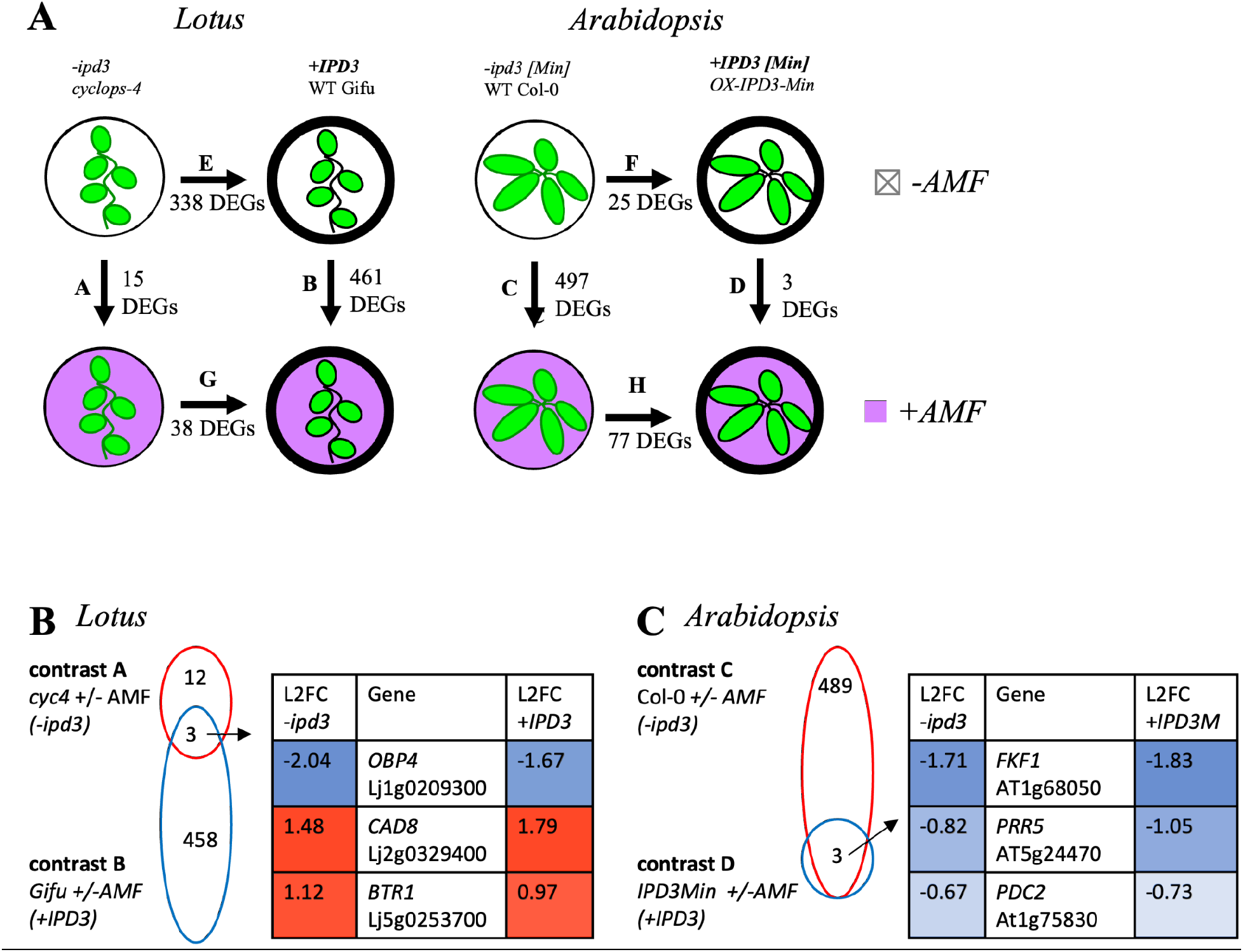
Differential transcription contrasts for *IPD3[Min]* expression with and without AMF treatment in two species. (A) Overview of all contrasts executed and number of DEGs detected. Contrast labels A-H in (B) and (C) and Figure 6 correspond to those noted in (A). DEGs reported for the *+IPD3Min* genotype in *Arabidopsis* are those attested in both of the individually conducted contrasts for the 2 transgenic lines 308 and 310. (B) Comparison of DEGs resulting from AMF treatment in *-ipd3* (*cyc4*) and *+IPD3* (Gifu) genotypes of *Lotus.* L2FC is Log2(fold change) of mean expression. (C) Comparison of DEGs resulting from AMF treatment in *-ipd3* (Col-0) and *+IPD3* (OX-*IPD3Min*) genotypes of *Arabidopsis.* Differential transcription is expressed as log2(fold change) (L2FC); more- positive numbers indicate higher expression and more-negative numbers indicate lower.

Next, we asked if the genes that remained responsive to AMF in *-ipd3 Lotus* would be conserved in *Arabidopsis.* To detect candidate genes that are part of the *Lotus* AMF host response but independent of *IPD3*, we looked for overlap between AMF-responsive DEGs for both genotypes (Figure 5B). Only 3 DEGs were shared: Lj1g0209300.1 (homolog of *OBF Binding Protein 4* (*LjOBP4*)); Lj2g0329400.1 (homolog of *Cinnamyl Alcohol Dehydrogenase 8* (*LjCAD8))*; and Lj5g0253700.1 (homolog of *Binding to TOMV RNA 1* (*LjBTR1*)). All of these genes were regulated in the same direction and at similar scale in both genotypes, suggesting they are elements of the native *Lotus* AMF response that are governed by an *IPD3-*independent mechanism. Despite *IPD3’s* effect on the scale of AMF response in *Lotus, IPD3^Min^* expression in *Arabidopsis* did not enable any new AMF-responsive DEGs, and the scale of response to AMF was much larger in the *-ipd3* Col-0 genotype than in *IPD3^Min^*(Figure 5C).

We then looked for overlaps in DEGs for AMF response between both genotypes of both species (Figure 6). For direct cross-species comparisons *Lotus* genes were labeled with their closest *Arabidopsis* homologs as recorded in genome annotations; total DEG count for AMF response of Gifu was slightly reduced due to exclusion of 33 genes without an annotated *Arabidopsis homolog* (Data S8). One of the 3 *IPD3-*independent genes characterized in Figure 5B as a member of the *IPD3*-independent *Lotus* AMF response, *OBP4* (*Lj1g0209300*; *At1g68050*), was also shared with the AMF response of Col-0 *Arabidopsis* (Figure 6A). *OBP4* is a transcription factor that negatively regulates lateral root and root hair development in response to nitrate and abscisic acid (Ramirez-Parra et al., 2017; Rymen et al., 2017; Xu and Cai, 2019). Interestingly, *OBP4* was upregulated in the AMF response of *Col-0 Arabidopsis*, but downregulated in *Lotus*. *OBP4* was not differentially regulated in *IPD3^Min^*plants.

**Figure 6:**
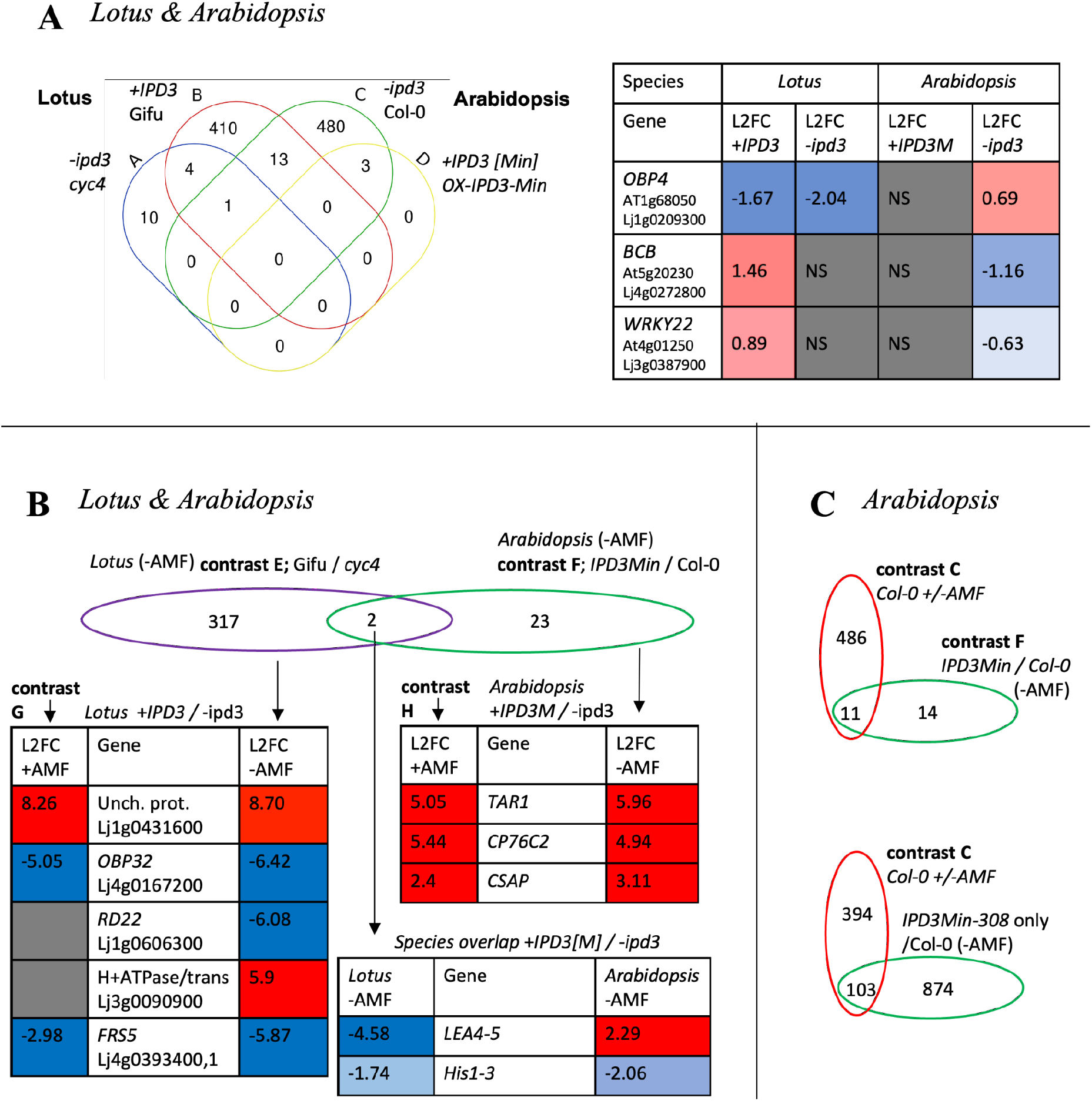
Cross-species comparison of AMF and *IPD3[Min]* responsive DEGs. (A) When the AMF-responsive DEGs within the respective +*IPD3* and *-ipd3* genotypes of *Lotus* and *Arabidopsis* are directly compared using annotated *Arabidopsis* homologs of *Lotus* genes, *OBP4* is shared by *-ipd3* Col-0 *Arabidopsis* with both *Lotus* genotypes, but oppositely regulated across species. Thirteen additional genes are shared by *-ipd3* Col-0 *Arabidopsis* and *+IPD3* Gifu *Lotus*, also with opposite regulation across species. (B) When the *IPD3-*responsive DEGs in non- AMF-treated *Lotus* and *Arabidopsis* are compared, *LEA4-5* and *His1-3* are shared across species, with *LEA4-5* being oppositely regulated. (C) When *IPD3-*responsive DEGs in non-AMF-treated *Arabidopsis* are compared to AMF- responsive DEGs in *-ipd3* Col-0 *Arabidopsis*, there is significant overlap. Eleven of 25 DEGs induced by *IPD3^Min^* in both transgenic lines 308 and 310 are also induced by AMF in *-ipd3* Col-0. 103 out of 497 AMF-responsive DEGs (21%) in *-ipd3* Col-0 are also responsive to *IPD3^Min^*in transgenic line 308, which has higher *IPD3^Min^* expression and a larger overall DEG set.

In addition to *OBP4*, 13 genes were differentially regulated in response to AMF in Gifu *Lotus* and Col-0 *Arabidopsis*, but not *cyclops-4 Lotus* or *IPD3^Min^ Arabidopsis* (Figure 6A, Data S8). In most of these genes, as for *OBP4*, the direction of regulation was reversed between species, including the next two most strongly regulated, multi-stress-responsive *Blue Copper Binding Protein* and pathogen response-related *WRKY22* (Figure 6A). Consistent with defense effects of AMF observed in the network analysis, *WRKY22*, which was downregulated in *Arabidopsis*, is implicated in systemic acquired resistance and is characterized as a positive regulator of the SA pathway in SA-JA defense crosstalk (Kloth et al., 2016).

We also compared the effect of *IPD3* presence and absence across the two species in the absence of AMF treatment. Two genes differentially transcribed in +*IPD3 Lotus* relative to *-ipd3 Lotus* were also differentially transcribed in *+IPD3^Min^ Arabidopsis* relative to *-ipd3* (Figure 6B). Of these, osmotic stress responsive *Late Embryogenesis Abundant 4-5* was strongly downregulated by the presence of *IPD3* in *Lotus* but upregulated by the presence of *IPD3^Min^* in *Arabidopsis*, while drought stress responsive *Histone H1-3* was downregulated in both species by the presence of respective *IPD3* versions.

Beyond specific genes, a shared and unexpected feature of the *+IPD3/-ipd3* contrast in both species was a pattern of many differentially expressed genes resulting from manipulation of IPD3 even in the absence of AMF (Figure 5A, Figure 6B). Many of the affected genes are related to biotic and abiotic stress. These included strong downregulation in *Lotus* of *Lj4g0393400.1* and *Lj1g0155200_LC.1,* both of which are homologs of *AtFRS5*, a gene involved in linking light perception to stress and pathogen defense via JA, and upregulation of *Lj4g0167200.1*, a homolog of the *AtJASSY* JA biosynthesis enzyme (Ma and Li, 2018; Guan et al., 2019). The top most strongly regulated gene that was dependent on the presence of *IPD3* in *Lotus* was *Lj1g0431600.1*; its homolog *Lj1g0070100.1* was also upregulated, with an average L2FC of 7.14. These genes are homologous to an uncharacterized MYB/SANT transcription factor in *Arabidopsis*, *At*2g24960.2. In *Arabidopsis*, *IPD3^Min^* expression also resulted in strong upregulation of genes related to JA pathogen response and auxin signaling including *Cytochrome P450 76C2* and *Tryptophan Aminotransferase Related 1* (Figure 6B)(Lorenzo et al., 2003; Stepanova et al., 2008). Notably, both of these genes were included in module E of the network analysis, which had the strongest correlation to quantitative *IPD3^Min^* expression (Data S6).

While the differential response to AMF was nearly eliminated in *-ipd3 Lotus*, 337 Differentially Expressed Genes (DEGs) respond to *IPD3* in the contrast of *+IPD3* and *-ipd3 Lotus* without AMF inoculation (Figure 5A). 135 of these DEGs resulting from knockout of *IPD3* in the absence of AMF treatment, were also found among DEGs resulting from AMF treatment of Gifu plants, accounting for 29% of the wildtype *Lotus* AMF response (Data S8). Thus, although AMF-responsive upregulation of genes by IPD3 is known to be essential for normal AM function, its deletion also appears to partially replicate the AM response under the conditions of this experiment. This suggests that *IPD3* in *Lotus* may act as a context-dependent transcriptional repressor of a portion of its own targets.

A similar pattern is present in *Arabidopsis*, where 11 of 25 genes (44%) differentially expressed in response to *IPD3^Min^*in the absence of AMF treatment were also induced by AMF treatment of Col-0 plants (Figure 6C). These genes included, for example, the *FKF1* circadian clock gene identified in Figure 5C as well as in the *IPD3-Min* responsive circadian clock module of the correlation network (Figures 4 & 5) (Data S8). In transgenic line 308 where higher *IPD3^Min^* expression corresponds to a larger absolute number of DEGs (Figure 5A, Figure S6), this effect is much larger. The no-AMF contrast of *IPD3^Min^-308* vs Col-0 recapitulates differential regulation of 103 of the 497 (21%) AMF-responsive DEGs in Col-0 (Figure 6C). These results confirm evidence in the correlation network that effects of *IPD3^Min^* expression in *Arabidopsis* overlap partially with the AMF response. They indicate that while the nature of the response may differ, expression of constitutively activated *IPD3* in *Arabidopsis* can activate the transcriptional response to AMF even in the symbiont’s absence, much as it does in AM host plants (Singh et al., 2014).

## Discussion

We successfully expressed *IPD3^Min^*, the DNA binding domain of the AM symbiosis-essential transcription factor IPD3, in the nonAM plant *Arabidopsis* which lost both the AM trait and *IPD3* 60-70 million years ago (Hohmann et al., 2015; Radhakrishnan et al., 2020). To understand the impact of *IPD3^Min^* on remaining genetic pathways and biotic interactions, we compared the response of *Arabidopsis* genotypes to AMF inoculation, as well as the AM host plant *Lotus* and its *ipd3* mutant *cyclops-4*. Our results indicate that despite the long intervening period as a nonAM plant and the further loss of related genes, expressing *IPD3^Min^* in *Arabidopsis* resulted in phenotypic and transcriptional effects. The preservation of molecular connections for *IPD3* raises the prospect of re-wiring nonAM plants to restore AM symbiosis, with applications in agriculture (French, 2017; Lynch, 2019). Although they are a phylogenetic minority, nonAM crops are responsible for almost half a billion metric tons of agricultural harvest every year (Hornstein, 2022). With reported yield increases from mycorrhizae of 20-100% in various host crops, significant economic advantages could result if these benefits can be conferred on nonAM crops (Eo and Eom, 2009; Pellegrino et al., 2015; Berruti et al., 2016). In 2021, such an increase in US canola alone would have resulted in an additional 250,000-1.25 million metric tons of harvest worth 175-870 million dollars (USDA NASS, 2022).

We also found evidence that *IPD3* may be subject to targeted regulation in *Arabidopsis*. Two full-length versions of *IPD3* transgenically expressed in *Arabidopsis* could not be detected at the protein level, while IPD3^Min^ was successfully expressed in roots and shoots (Figure 2, S1). This suggests *IPD3* may be subject to silencing or degradation specific to the N-terminal portion excluded from IPD3^Min^. In prior work, we attempted to express *IPD3^Mt^*and *IPD3^S50D^* in the oilseed crop *Camelina sativa*, and observed low expression levels in multiple independent lines, consistent with evidence in the present study for silencing of these constructs in *Arabidopsis* (Hornstein 2022). We have also observed potentially deleterious effects for *IPD3^S50D^* in *Camelina.* T1 transformants were repeatedly lost to severe fungal infections, and later generations produced unusual dwarfed growth phenotypes at nonmendelian ratios (Hornstein and Sederoff, unpublished). It is possible that both the present study and past work in *Camelina* were biased by survivorship of T1 individuals which avoid a deleterious effect due to silencing.

Mechanisms for functional AM go beyond the CSP to essential plant functions including lipid biosynthesis, membrane and vesicle functions, nutrient transport, and defense (Wang et al 2012; Behie and Bidochka, 2014; Luginbuehl et al., 2017). Although symbiosis-specific genes that effect these functions in AM hosts have been lost in nonAM species, most belong to families with non-symbiosis-specific members that have similar molecular functions. In our experiment we observed *IPD3-*independent upregulation of lipid and 2-MAG biosynthesis genes in response to AMF in *Arabidopsis* (Figure 3, Figure 4). Activation of 2-MAG biosynthesis via *RAM1* and *RAM2* is well-characterized as an essential feature of AM symbiosis that strictly depends on *IPD3* in host species (Wang et al., 2012; Gobbato et al., 2013; Pimprikar and Gutjahr, 2018). In our results, 2-MAG-related transcription appeared to occur in response to AMF, but by action of related genes which are not symbiosis-specific (Yang et al., 2010; Yang et al., 2012b). In the context of engineering AM as an agriculturally useful trait, this raises the interesting possibility that non-symbiosis-specific genes retained in nonAM plants can be recruited to perform symbiotic functions (Hornstein and Sederoff, in preparation).

Figure 7 lays out the main findings of this study in relation to the canonical knowledge of *IPD3*’s role in AM. In *Arabidopsis, IPD3^Min^* plants showed a >99% reduced transcriptional response to AMF compared to wild type (Figure 5). This is in part because *IPD3^Min^* expression places plants in an AMF-exposure-like transcriptional state prior to actual exposure to the fungus (Figure 6C, S4). This is reminiscent of results in symbiosis models where expression of phosphomimic *IPD3* leads to a constitutive symbiotic response in the absence of biotic stimulus (Singh et al., 2014; Gobbato, 2015). Our results suggest that although the nature of the transcriptional response may differ, *IPD3^Min^* acts analogously in *Arabidopsis* by activating the native AMF response even in the absence of the fungus. In *Lotus*, surprisingly, we found that - *ipd3* <u>knockout</u> was associated with a similar priming effect. The AMF response in this *Lotus* genotype is ∼90% reduced in our experiment, consistent with canonical understanding of *IPD3* (Yano et al., 2008; Pimprikar et al., 2016). However, in our experiment this was partly because a contingent of genes in uninoculated plants is regulated similarly upon *ipd3* knockout and AMF exposure (Figure 5, Figure S5). This suggests IPD3 may have an unrecognized function as a direct or indirect repressor of some of its own targets.

**Figure 7:**
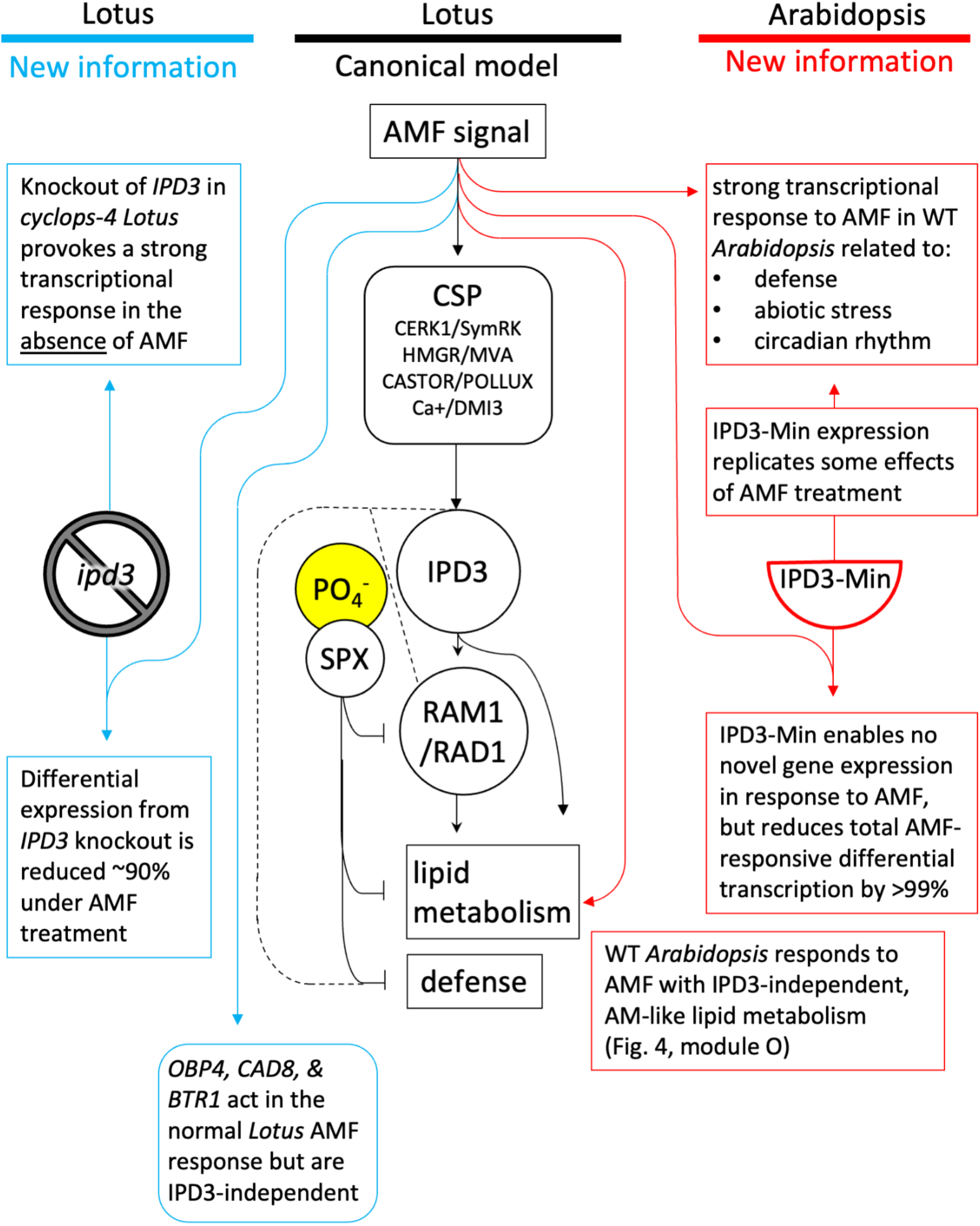
Novel roles for *IPD3* and responses to AMF in *Arabidopsis* and *Lotus*. In *Lotus*, we add to the current understanding of *IPD3* by showing that its knockout has a large effect even in the absence of AMF, and that the reduction in overall scale of AMF response in the *cyclops-4* genotype is due to a surprising partial replication of AMF-exposure-like transcription. In *Arabidopsis* we find evidence that AM-related lipid metabolism is altered by AMF independently of *IPD3* and despite the plant’s nonAM status. Expression of *IPD3^Min^* vastly reduced the amount of AMF-responsive differential expression in *Arabidopsis*, but not because *IPD3^Min^* has little effect. *IPD3^Min^*expression results in many differentially expressed genes in the absence of AMF, and the response to AMF in transgenic plants is small at least in part because many of the DEGs acting in the wild type *Arabidopsis* AMF response are similarly regulated in *IPD3^Min^*transgenics even before AMF treatment is applied. Genes affected by *IPD3^Min^*in *Arabidopsis* relate to functions including abiotic stress, pathogen defense, and circadian rhythm.

The striking inverse effects in *Lotus* and *Arabidopsis* also highlight two limitations of our results for drawing comparisons between the two species and with prior work. First, the *IPD3* genotypes across the two species are not identical. The native *LjIPD3/CYCLOPS* gene in *Lotus* encodes a protein product that undergoes extensive protein-protein interactions and requires phosphorylation for activation, while the IPD3^Min^ protein we used in *Arabidopsis* is constitutively active (Yu et al., 2014; Singh et al., 2014; Pimprikar et al., 2016; Jin et al., 2016; Jin et al., 2018). Second, the effects we observed for AMF interaction may be specific to the early timepoint sampled in this study (48h). We selected an early time point for the whole experiment because *Arabidopsis* cannot sustain AMF in monoxenic culture, but most studies allow interaction with AMF on the scale of weeks to establish mature colonization; earlier stages can involve rapidly shifting responses not seen later (Siciliano et al., 2007; Genre et al., 2008; Gutjahr et al., 2009; Handa et al., 2015; Nanjareddy et al., 2017; Prihatna et al., 2018). An idealized experiment might use *cyclops-4 Lotus* rescued by expression of *IPD3^Min^*, and a nurse- pot colonization system to support *Arabidopsis* AM colonization (Veiga et al., 2013; Fernández et al., 2019).

One area where *IPD3^Min^* expression duplicates the effect of AMF treatment is downregulation of circadian clock genes that result in delayed flowering (Figure 4) (Sawa et al., 2007; Para et al., 2007; Niinuma et al., 2008; Shim et al., 2017). This gene regulation may explain the delayed flowering phenotype in *IPD3^Min^* transgenic *Arabidopsis* seen in our experiment if the root transcriptome effect extends to aboveground tissue (Figure 1; Figure S1). AM symbiosis affects flowering time in host plants, and the circadian clock is implicated in maintenance of the symbiosis itself as well as AM-mediated abiotic stress resistance (Hernandez and Allen, 2013; Lee et al., 2019; Bennett and Meek, 2020; Liu et al., 2022). In *Arabidopsis*, circadian rhythm genes are also known to act in microbiome construction, pathogen defense, development, and abiotic stress (Lee et al., 2005; Nakamichi et al., 2016; Newman et al., 2022; Xu et al., 2022; Singh, 2022a; Singh, 2022b). Remarkably, in mosses, the sole group of nonAM plants that retain CSP genes, *IPD3* specifically has been shown to mediate a stress-responsive reproductive transition (Kleist et al., 2022).

In uninoculated plants, two DEGs result from the respective *+IPD3/-ipd3* contrasts in both *Arabidopsis* and *Lotus*: *LEA4-5* and *HIS1-3* (Figure 6B). Both of these genes are associated with response to abiotic stress including drought, heat, and cold (Dalal et al., 2009; Olvera-Carrillo et al., 2010; Rutowicz et al., 2015). *HIS1-3* is downregulated in the *+IPD3* genotype of both species, and *LEA4-5* is upregulated in *Arabidopsis* but downregulated in *Lotus* (Figure 6). A molecular function in stress response would subject *IPD3* to selection on factors unrelated to its role in AM symbiosis, and might help explain both the reasons for this gene’s loss in nonAM species, and its presence in charophyte ancestors of land plants long before the existence of AM symbiosis (Delaux et al., 2015; Delaux and Schornack, 2021).

We also identified AMF and *IPD3-Min*-responsive downregulation of *Arabidopsis* genes involved in salicylic acid-mediated defense (Figure 4C). Reduced transcription of *WRKY70*, *WRKY54*, and *PR5* by *IPD3^Min^* in the absence of AMF suggests that plants expressing *IPD3^Min^* have reduced baseline levels of SA-mediated defense to biotrophic pathogens (Glazebrook, 2005; Blanco et al., 2005; Li et al., 2006; Yang et al., 2012a). Early AM colonization of host plants can involve a transient increased defense response followed by reduction, while *Arabidopsis* shows an early symbiosis-like response to AMF, followed by a strong defensive response under forced long-term interaction (Giovannetti et al., 2015) (Fernández et al., 2019; Cosme et al., 2021). While AM can confer resistance to pathogens and insects, genes acting in AM also enable infection by some pathogens (Wang et al., 2012; Siebers et al., 2016; Ried et al., 2019; Chen et al., 2021a; Dey and Ghosh, 2022). The full mechanisms for such effects are not known, but comport with our evidence for perturbation of *Arabidopsis* defenses by a nominally AM-specific gene.

Recently, significant portions of the AMF genetic network were revealed to be under control of the phosphate-stress-responsive PHR-SPX system, which is conserved in *Arabidopsis* (Shi et al., 2021; Shi et al., 2022). In *Arabidopsis*, PHR also regulates the microbiome by reducing plant defenses when phosphate is low, resulting in recruitment of beneficial microbes (Finkel et al 2019, Martin-Rivilla et al. 2019, Castrillo et al 2017,Cho, Kang, and Kim 2013; Kotchoni and Gachomo 2006; Dangl 1998). Given knowledge that both AM and non-AM microbial relations are part of the PHR-regulated phosphate response network, it is possible that the *IPD3^Min^* effect in our experiment relates to conserved, non-AM-specific points of crosstalk with defense and symbiosis (Figure 7).

Regardless of whether plants are formally labeled non-hosts in isolation, in natural settings they continue to interact with ubiquitous AM fungi and can display partial and transitional phenotypes (Ma et al., 2018; Cosme et al., 2018). It is not surprising that an ancient, conserved trait is interconnected to other aspects of plant life. Our results show that despite the loss of genes essential for AM symbiosis, substantial connections to this trait remain in place in *Arabidopsis*, and can be highlighted, even re-activated by expression of *IPD3^Min^*. The transcriptional effects we identified suggest specific targets for follow-up studies to directly assess pleiotropic effects, including pathogen sensitivity and abiotic stress resilience.

## Methods

Please see supplemental experimental procedures for additional detail on all sections.

### Generation of transgenic *Arabidopsis*

The coding sequences of MtIPD3, S50D-IPD3, and IPD3-Min (Genbank EF569224.1; Yano et al. 2008, Singh et al 2014) were synthesized (Integrated DNA Technologies, Research Triangle Park, NC) and assembled by Hi-Fi assembly (New England Biolabs, Ipswich, MA) into a modified pCAMBIA0380 expression construct (Genbank AF234290.1) under control of the Arabidopsis Ubiquitin 10 promoter (Ivanov and Harrison 2014). *35S:mCherry* amplified from pC-GW-mCherry (Genbank KP826771.1) (Dalal et al. 2015) was the selection marker.

Arabidopsis were transformed as described by Davis et al. (2009). Seeds were screened by fluorescence and PCR during segregation and lines were brought to homozygosity.

### Protein analysis

Protein was extracted from roots and shoots of 5-week-old seedlings grown on ½ MS plates. For untargeted shotgun proteomics, 5 samples per line and tissue type were ground and lysed in SDT buffer, then prepared as described in Wiśniewski et al. (2009) before 1D-LC-MS/MS using an UltiMate^TM^ 3000 RSLCnano Liquid Chromatograph and Orbitrap Eclipse Tribrid mass spectrometer (Thermo Fisher) as described in Mordant and Kleiner (2021). Mass spectra were searched against a database of Col-0 *A. thaliana* proteins (Uniprot:UP000006548) using the SEQUEST HT algorithm in Proteome Discoverer. Protein abundance was quantified as normalized spectral abundance factor (NSAF) in Microsoft Excel (Zybailov et al 2006).

For analysis of specific protein size ranges corresponding to bands observed on blots or predicted protein length, we used in-gel digestion of excised SDS-PAGE prior to LC-MS/MS analysis (GeLC-MS/MS). Only 1 sample per construct was used for GeLC-MS/MS. 30-40 µg of total protein in lysate generated for untargeted proteomics was denatured by heating with Laemmli buffer, then run on an SDS-PAGE 12% separating gel with 5% stacking gel. Excised bands were processed according to Shevchenko et al (2006), and 10 uL of peptide mixture was injected to LC as described above, connected to an Orbitrap Exploris 480 mass spectrometer (Thermo-Fisher) for MS/MS with the same settings used for untargeted proteomics.

For Western blotting, frozen tissue was hand-ground with extraction buffer, centrifuged, and the supernatant denatured by heating in LDS sample buffer. Protein was run on a 12% Tris-Glycine acrylamide gel, then transferred to PVDF membranes and blocked in TBS+2% BSA with 4uL/mL Tween-20, then incubated with 1:1000 dilution of custom rabbit anti-IPD3 polyclonal antibody (Genscript, China) followed by 1:2,500 donkey anti-rabbit AlexaFluor 488- conjugated secondary antibodies (Thermo Fisher, Waltham, MA) and imaged on a GelDoc SR (Bio-Rad, Hercules, CA).

### Growth phenotyping

Plants were grown in the NCSU phytotron under long day conditions in 8 oz pots filled with SunGro propagation mix (Sungro, Agawam, MA). Pots were hand-watered with deionized water. Seeds were harvested from dried mature plants and hand-cleaned before weighing. The seed yield experiment was repeated 3 times with n=9-15. The flowering time experiment was performed once with n=14 for line 312 and n=15 for all other lines, with plants censused daily for onset of bolting.

### Transcriptome experiment

Seeds were sterilized and grown for 5 weeks under long-day conditions (16 hours light/8 hours dark, 21/18C) on sterile petri dishes containing either 1/2MS or low-nutrient MS media (Phytotech Labs, Lenexa, KS). 48 hours prior to sample collection, ∼200 germinated *R. irregularis* spores (Premier Tech, Canada) or a water mock inoculum were added to the roots of AMF-treated plants. Each replicate consisted of the pooled roots of 5 seedlings from the same plate.

RNA was extracted with the Purelink RNA Mini kit and treated with Turbo DNAse (Invitrogen, Waltham, MA), then sequenced by DNBSEQ at BGI Group (China) using strand-specific, poly-A enrichment to obtain 100 bp paired-end reads. *Lotus* and *Arabidopsis* alignments were performed using BBSplit, a multi-reference aligner, to separate plant and fungal reads (Bushnell 2014). Reference genomes used were Araport 11 for *Arabidopsis* (TAIR), the Joint Genomics Institute assembly for *R. irregularis* (Genbank: GCA_000439145.3), the Gifu V1.2 assembly for *Lotus* (GCA_012489685.2), the *IPD3-Min* transformation construct, and the *LjCYCLOPS* sequence. Reads were counted at the transcript level using featureCounts (Liao, Smyth, and Shi 2014).

Gene expression networks were constructed using WGCNA v1.69 (Langfelder and Horvath, 2008; 2012) with soft-threshold power of 16 (*Arabidopsis*) or 24 (*Lotus*). Correlation coefficients were calculated between the eigengenes of each module and treatment variables to identify significant module-trait relationships, with p<0.05. Gene Ontology enrichment and hierarchical term clustering was performed with PANTHER (Mi et al. 2013). Terms were filtered by specificity and significance, then subjected to semantic similarity clustering in reviGO (Supek et al 2011).

Differential expression analysis was performed in R using the edgeR package (Liao, Smyth, and Shi 2014; Robinson, McCarthy, and Smyth 2010). The estimateGLMCommonDisp function with FDR-adjusted p-value <0.05 was used to test for differential expression. Cross-comparison of gene lists was done in Excel.

## Data Availability

Transcriptome data is publicly available via GEO accession number GSE225213. The mass spectrometry proteomics data have been deposited to the ProteomeXchange Consortium via the PRIDE partner repository with the dataset identifier PXD040665 (prepublication access at www.ebi.ac.uk/pride/login with Username: and Password: CVDhdrfh)

## Acknowledgements

We thank Swathi Barampuram, Matthew Occena, and Asa Budnick for assistance with laboratory work in this study. We thank Caroline Gutjahr and Makoto Hayashi for use of the *cyclops-4 Lotus* mutant. This work was supported by the Novo Nordisk Foundation InRoot project under award No. NNF19SA0059362 (H.S. & M.K.) and the Department of Energy, DE- SC0018269 (H.S., M.C., E.D.H.). Graduate student fellowships from the National Science Foundation (NRT-INFEWS #1828820) and NCSU Provost’s Doctoral Fellowship provided support for E.H. and NIH Molecular Biotechnology Training Grant **#** 1T32GM133366-01 Fellowship to M.C. and a UsDoEd GAANN fellowship (P200A210002) to M.F.. The LC-MS/MS measurements of protein samples were made using equipment in the Molecular Education, Technology, and Research Innovation Center (METRIC) at North Carolina State University.

## Author Contributions

E.D.H. and H.S. designed the research; E.D.H. and S.V. performed the research; B.E., S.V., and M.K. contributed analytical tools; M.C., M.F., S.V., and E.D.H. analyzed the data; EDH and HS wrote the paper with contributions from all authors.

## Supplemental experimental procedures

### Assembly of transgenic constructs

Primer pairs for cloning and screening are shown in the table contained within this section. The coding sequence of *Medicago truncatula IPD3* (Yano et al. 2008), (Genbank accession EF569224.1) was synthesized including restriction sites for *BamHI* (5’) and *SphI* (3’) (Integrated DNA Technologies, Research Triangle Park, NC) and assembled into pUC19 (Addgene plasmid #50005) using respective enzymes and T4 ligase (NEB, Ipswich, MA), followed by transformation and selection with carbenicillin. The *S50D-IPD3* coding sequence was constructed by restriction-ligation of pUC19-*MtIPD3* with a synthesized fragment containing the first 781 bases of *IPD3* including the S50D mutation (IDT, Research Triangle Park, NC) via *Pst1* and *BamHI* (NEB, Ipswich, MA).

Plant expression constructs were prepared in the pCAMBIA0380 (Genbank AF234290.1) backbone by seamless assembly with the NEB Hi-Fi cloning kit (NEB, Ipswich, MA). Amplifications for Gibson assembly used Superfi Platinum II polymerase (Invitrogen, Waltham, MA). Assembly fragments were digested with *DpnI* and gel-purified with the NEB Monarch gel extraction kit (NEB, Ipswich, MA). Plasmids were transformed into *E. coli*, recovered and plated onto LB with appropriate antibiotics.

Colonies screened by direct PCR were used to inoculate liquid culture and plasmid was purified from 3- 100 mL of saturated culture using Qiaprep Spin Miniprep or Zymo Midiprep kits and Sanger-sequenced for confirmation, including plasmids produced as intermediate cloning steps (Qiagen, Germany; Zymo, Irvine, CA).

First, the 2X35S:mCherry marker sequence amplified from pC-GW-mCherry (Genbank KP826771.1) (Dalal et al. 2015) was assembled into the empty marker site of pCAMBIA0380 to produce pC0380-MC using primer pairs 1 and 2. Next, a synthetic DNA fragment containing the 35S promoter, *Nopaline Synthase* terminator (*tNOS*), and *Arabidopsis Heat Shock Protein* terminator (*tAtHSP*), was then assembled into the multiple cloning site (MCS) of pC0380-mCherry linearized with primer pair 3 to produce pC0380-MC-35S (Nagaya et al. 2010). The 2 kb *Arabidopis Ubiquitin 10* promoter (*pAtUBQ10*) was amplified from Col-0 *Arabidopsis* genomic DNA with primer pair 4 and assembled in pC0380-MC-35S linearized with primer pair 5 to produce pC0380-MC-UBQ, in which *pAtUBQ10* replaces *p35S* for the gene of interest (Ivanov and Harrison 2014). The purpose of the *tNOS:AtHSP* double terminator was to increase expression and the purpose of replacing *p35S* with *pATUBQ10* was to provide a promoter with expression document in multiple root tissues as described by Nagaya et al and Ivanov and Harrison, respectively (Ivanov and Harrison 2014; Nagaya et al. 2010).

*MtIPD3* or *S50D-IPD3* coding sequences were amplified from respective pUC19 cloning plasmids with primer pair 6 and assembled in position following *pAtUBQ10* in pC0380-MC-UBQ linearized with primer pair 7. *IPD3-Min* was cloned by amplifying the N-terminal DNA-binding domain (aa 254-513) as described in Singh et al (2014) from *pUC19-MtIPD3* with addition of a 5’ start codon using primer pair 8, and assembled in pC0380-MC-UBQ.

### Plant transformation

Col-0 *Arabidopsis* were transformed by the direct-dip protocol as described by Davis et al. (2009).

Briefly, *Agrobacterium tumefaciens* GV3101 transformed with expression constructs was grown to saturation in yeast extract-beef medium with addition of 2.5% sucrose and appropriate antibiotics and screened by colony PCR with primer pair 9 or 10 to confirm presence of the trangene. An additional 2.5% sucrose and 300 uL/L Silwet L-277 (Phytotech Labs, Lenexa, KS) were added to bacterial cultures and flowering *Arabidopsis* were dipped directly into this mixture; bacteria were not resuspended in transformation buffer and vacuum was not used. Plants were covered in the dark for 24 hours then returned to growing conditions; transformation was repeated 3 times, 4-6 days apart.

T1 *Arabidopsis* seed were screened by mCherry fluorescence, and sequence of the transgene was confirmed by CTAB DNA extraction, PCR with primer pairs 9 and/or 10, and Sanger sequencing (Porebski, Bailey, and Baum 1997). RNA was extracted from leaves using the Invitrogen Purelink RNA Mini kit (Invitrogen, Waltham, MA), treated with the Turbo DNA-free DNAse kit (Invitrogen, Waltham, MA) and screened for expression of the transgene by PCR of cDNA generated with the Takara EcoDry cDNA kit (Takara Bio, Japan). Lines were brought to homozygosity over 3 generations, with PCR and Sanger sequence confirmation of the transgene in each generation.

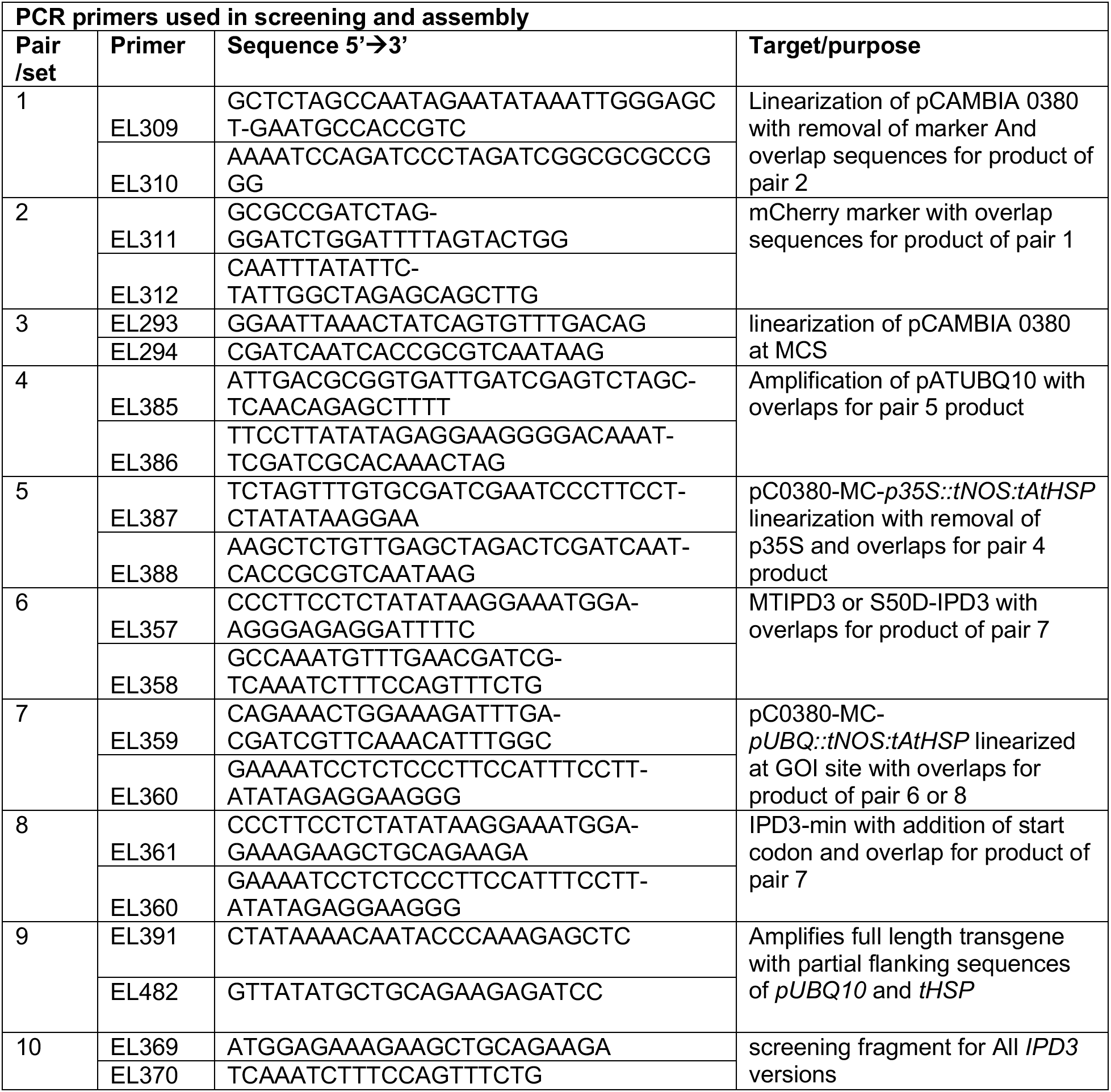

### Protein analysis

For Western blotting, tissue was frozen in liquid nitrogen and ground in a mortar and pestle. Extraction buffer consisting of 50 mM Tris-HCL pH 8, 150 mM NaCl, .1 mM Brij-35, 1 mM EDTA, 2.5 mM DTT, 2% SDS w/v, 10% glycerol v/v, and freshly added 2% 2-mercaptoethanol v/v and 1.5% Sigma plant protease inhibitor cocktail (Sigma-Aldrich, Burlington, MA) was added to the frozen tissue and ground until thawed. Samples were centrifuged at 4C for 10 minutes, and the supernatant was diluted 1:1 in NuPage LDS sample buffer (Invitrogen, Waltham, MA) with an additional 2% v/v 2-mercaptoethanol and heated at 95C for 10 minutes. 45 uL of sample was run on a Novex Wedgewell 12% Tris-Glycine (Invitrogen, Waltham, MA) acrylamide gel in SDS running buffer for approximately 25 minutes at 225 mV.

Protein was transferred to PVDF membranes using an iBlot (Thermo Fisher, Waltham, MA). Membranes were blocked in Tris-buffered saline plus 2% BSA and 4uL/mL Tween-20 for 1 hour, then incubated for at least 16 hours at 4C with 1:1000 dilution of custom rabbit anti-IPD3 peptide primary polyclonal antibody (Genscript, China) in blocking buffer. Membranes were washed 3 times in blocking buffer, then incubated at RT in TBS-T with 1:2,500 dilution of donkey anti-rabbit Alexa Fluor 488-conjugated fluorescent secondary antibodies (Thermo Fisher, Waltham, MA) for 2-4 hours. Blots were imaged on a GelDoc SR (Bio-Rad, Hercules, CA).

For shotgun proteomics, ground leaf and root tissue was used for protein extraction. Five biological replicates were included for each line. We used 200-300 mg of hand-ground tissue from each sample for protein extraction in 1 ml of SDT lysis buffer [4% (w/v) SDS, 100 mM Tris-HCl pH 7.6, 0.1 M DTT]. We lysed the ground tissue by bead-beating in lysing matrix E tubes (MP Biomedicals) with a Bead Ruptor Elite (Omni International) for 5 cycles of 45 sec at 6.45 m/s with 1 min dwell time between cycles; followed by heating to 95°C for 10 min. The lysates were centrifuged for 5 min at 21,000 x g to remove cell debris. Supernatant was used for purification and digestion using the filter-aided sample preparation (FASP) protocol described by Wisniewski et al. (2009). All centrifugations mentioned below were performed at 14,000 x g. Samples were loaded onto 10 kDa MWCO 500 μl centrifugal filters (VWR International) by combining 60 μl of lysate with 400 μl of Urea solution (8 M urea in 0.1 M Tris/HCl pH 8.5) and centrifuging for 20 min. This step was repeated once to load a total of 120 ul of lysate. Filters were washed once by applying 200 μl of urea solution followed by 20 min of centrifugation to remove any remaining SDS. 100 μl IAA solution (0.05 M iodoacetamide in Urea solution) was added to filters for a 20 min incubation at room temperature followed by centrifugation for 20 min. The filters were washed three times with 100 uL of urea solution and 20 min centrifugations, followed by a buffer exchange to ABC (50 mM Ammonium Bicarbonate). Buffer exchange was accomplished by three cycles of adding 100 μl of ABC and centrifuging for 20 min. Tryptic digestion was performed by adding 1 μg of MS grade trypsin (Thermo Scientific Pierce, Rockford, IL, USA) in 40 μl of ABC to each filter and incubating for 16 hours in a wet chamber at 37°C. Tryptic peptides were eluted by adding 50 μl of 0.5 M NaCl and centrifuging for 20 min. Peptide concentrations were determined with the Pierce Micro BCA assay (Thermo Scientific) following the manufacturer’s instructions.

For analysis of specific protein size ranges we used SDS-PAGE followed by in-gel digestion of specific bands. We used the proteomics lysate in SDT buffer generated for the shotgun proteomic approach also for SDS-PAGE. Only one replicate of each transgenic line was used for SDS-PAGE. We mixed lysate containing 30-40 µg of protein with Laemmli buffer, and heated to 95°C for 5 min prior to loading. SDS-PAGE was done using a 12% separating gel with a 5% stacking gel that was run at 80 V for 30 min and then 120V for 1 hour and 15 min. The gel was fixed with 40% EtOH and 10% acetic acid for 30 min followed by staining with QC Colloidal Coomassie stain (Bio-Rad) overnight. Gel pieces corresponding to target protein sizes were excised and in-gel digestion was performed according to Shevchenko et al. (2006). Gel pieces were destained in 40 mM ammonium bicarbonate buffer with 50% acetonitrile. Reduction was performed using 20 mM DTT for 30 min at 56°C and alkylation was performed using 55 mM iodoacetamide for 20 min at room temperature. In-gel digestion was performed overnight using a 0.02 µg/µl trypsin solution. Elution of peptides was done by addition of 100% acetonitrile followed by a second elution using 50% acetonitrile and 5% formic acid. The peptide mixture was dried in a speedvac and rehydrated using 10 µl 2% formic acid.

Shotgun proteomics samples were analyzed by 1D-LC-MS/MS as described in Mordant and Kleiner (2021). The samples were blocked by treatment. Nontransgenic control samples were run first to avoid any false detections due to minimal amounts of carryover of peptides from the transgenic plant samples to control samples. We loaded of 1.4 μg peptide of each sample with an UltiMate^TM^ 3000 RSLCnano Liquid Chromatograph (Thermo Fisher Scientific) in loading solvent A (2% acetonitrile, 0.05% trifluoroacetic acid) onto a 5 mm, 300 µm ID C18 Acclaim® PepMap100 pre-column (Thermo Fisher Scientific). Elution pre-column and separation of peptides on the analytical column (75 cm x 75 µm analytical EASY-Spray column packed with PepMap RSLC C18, 2 µm material, Thermo Fisher Scientific; heated to 60 °C) was achieved using a 140 min gradient going from 95 % buffer A (0.1 % formic acid) to 31 % buffer B (0.1 % formic acid, 80 % acetonitrile) in 102 min, then to 50 % B in 18 min, to 99 % B in 1 min and ending with 99 % B. The analytical column was connected to an Orbitrap Eclipse Tribrid mass spectrometer (Thermo Fisher Scientific) via an Easy-Spray source. Eluting peptides were ionized via electrospray ionization (ESI). Carryover was reduced by two wash runs (injection of 20 µl acetonitrile, 99 % eluent B) between sample blocks. MS1 spectra were acquired in the Orbitrap by performing a full MS scan at a resolution of 60,000 on a 380 to 1600 m/z window. MS2 spectra were acquired using a data dependent approach by selecting for fragmentation the 15 most abundant ions from the precursor MS1 spectra. We used a normalized collision energy of 27 for HCD in the ion-routing multipole to generate the peptide fragments for MS2 spectra. Other settings of the data-dependent acquisition included: a maximum injection time of 50 ms, a dynamic exclusion of 25 sec and exclusion of ions of +1 charge state from fragmentation. About 50,000 - 60,000 MS/MS spectra were acquired per sample.

For measurement of gel bands after in-gel digest, 10 µl of peptide mixture was injected using the same Liquid Chromatograph and protocol as above. The analytical column was connected to an Orbitrap Exploris 480 mass spectrometer (Thermo-Fisher Scientific) with the same MS1 and MS2 parameters used on the Eclipse. About 95,000 MS/MS spectra were acquired per sample.

### Protein identification and quantification

A database containing all protein sequences from *A. thaliana* cv. Col-0 (Uniprot:UP000006548), as well as the IPD3 vector sequences was used. Sequences of common laboratory contaminants were included by appending the cRAP protein sequence database (http://www.thegpm.org/crap/). The final database contained 39,446 protein sequences and is included in the PRIDE submission (see data access statement) in fasta format. Searches of the MS/MS spectra against this database were performed with the Sequest HT node in Proteome Discoverer version 2.3.0.523 (Thermo Fisher Scientific) as described in Mordant and Kleiner (2021). Peptide false discovery rate (FDR) was calculated using the Percolator node in Proteome Discoverer and only peptides identified at a 5% FDR were retained for protein identification. Proteins were inferred from peptide identifications using the Protein-FDR Validator node in Proteome Discoverer with a target FDR of 5%. The data were normalized by calculating normalized spectral abundance factors (NSAFs) according to Zybailov et al. (2006) and multiplied by 100 to represent the relative protein abundance as a percentage.

### Growth phenotyping

Plants were grown in the NCSU phytotron under long day conditions in 8 oz pots filled with SunGro propagation mix (Sungro, Agawam, MA). Pots were hand-watered daily with deionized water and were not fertilized. Plants were censused daily for onset of bolting, onset of flowering, and duration of seeding. Mature plants were dried for 1 week, then seeds were manually harvested and hand-cleaned to remove chaff before weighing.

### Anthocyanin extraction

To check for anthocyanins and carotenoids in visibly red roots of transgenic plants, roots of mature soil-grown plants were briefly washed, then frozen with liquid nitrogen and hand-ground. 600 uL of pure methanol, 60% methanol/water v/v + 1% HCl, or 80% acetone was added to ∼100 mg of ground tissue and further ground with a plastic pestle in a 1.7 mL microtube, then centrifuged for 10 minutes at 16,000 x g, and supernatant harvested. Absorbance was measured at 10 nm intervals using a Synergy HT plate reader (Biotek, Winooski, VT).

### Transcriptome experiment preparation

Plants were grown on sterile petri dishes containing 50 mL of either 1/2MS or low-nutrient MS (LN) media. LN media was prepared from 20X macronutrient solution and MS micronutrient solution (Phytotech Labs, Lenexa, KS) as described in the table below. Plants were separated from growing medium by a 30 uM nylon mesh (Genesee, RTP NC) to prevent roots from growing into the media. Plates were covered with black cardboard sleeves to prevent light exposure of roots, and held at a 60 degree angle during growth in a growth chamber under long-day conditions (16 hours light/8 hours dark). *Lotus* seeds were treated identically to *Arabidopsis* with the exception of being manually scarified on 300-grit sandpaper prior to sterilization.

| Ingredients and macronutrient concentrations of nutrient media used in this study along with preprepared micronutrients at standard ½ MS concentration. |  |  |  |  |
| --- | --- | --- | --- | --- |
| Reagent | Macronutrient | mM in ½ MS | mM in LNMS | LN as % of 1/2MS |
| KH<sub>2</sub>PO<sub>4</sub> | -- | 0.625 | 0 | -- |
| K<sub>2</sub>HPO<sub>4</sub> | -- | 0 | 0.0625 | -- |
| NH<sub>4</sub>NO<sub>3</sub> | -- | 10.3 | 2 | -- |
| KNO<sub>3</sub> | -- | 9 | 2 | -- |
| MgSO<sub>4</sub>*7H<sub>2</sub>O | -- | 0.75 | 0.75 | -- |
| CaCl<sub>2</sub> | -- | 1.5 | 1.5 | -- |
| -- | NH<sub>4</sub> | 10.3 | 0.2 | 0.97% |
| -- | NO<sub>3</sub> | 19.3 | 0.1 | 1.04% |
| -- | total N | 29.6 | 0.3 | 1.01% |
| -- | PO<sub>4</sub> | 0.625 | 0.003 | 0.50% |
| -- | K | 9.625 | 0.11 | 1.10% |

500 uL of AMF inoculum containing ∼200 aseptic *R. irregularis* spores (Premier Tech, Canada) germinated at 26C for 1 week prior to use was pipetted onto the roots of each plant. Control plants were mock-inoculated with sterile water. Plants were allowed to sit horizontally for 2 hours after inoculation, then returned to the growth chamber for the remainder of the 48 hours prior to collection. All treatments and tissue collection were completed between 4 and 6 hours from the start of the light period, and roots were immediately frozen in liquid nitrogen. Each replicate consisted of the pooled roots of 5 seedlings from the same plate.

RNA was extracted and DNAse treated as described previously, then sequenced by BGI Group (China). RNA libraries were prepared at BGI using a strand-specific, poly-A enrichment method and then sequenced on the DNBSEQ platform to obtain 100 bp paired-end reads. Raw reads were filtered for adaptor contamination and low-quality sequences using SOAPnuke (Chen et al. 2018) and remaining sequences were evaluated using FASTQC (Andrews 2010) to ensure only high-quality read pairs (Q>30) were used for downstream analysis.

Read alignment was performed using BBSplit, an aligner designed for metagenomics within the BBTools bioinformatics toolkit (Bushnell 2014), in order to align three reference genomes simultaneously. Reference genomes used for *Arabidopsis* were the Araport 11 assembly for Arabidopsis (TAIR), the Joint Genomics Institute genome assembly for *R. irregularis* (Genbank: GCA_000439145.3), and a synthetic reference genome containing the T-DNA sequence as well as the *mCherry* selection marker and *AtUBQ10* and *2X35S* promoter sequences. *Lotus* assembly used the Gifu V1.2 assembly (GCA_012489685.2) and a synthetic genome containing the *CYCLOPS/LjIPD3* sequence. Reads were assigned to one of three reference genomes based on the best alignment score and ambiguous reads were discarded. Reads that aligned to an annotated feature in the genome were summarized using featureCounts (Liao, Smyth, and Shi 2014). Genes with zero counts or with overall low abundance in all samples (<10 read counts) were subsequently filtered prior to downstream analysis.

Gene expression networks were constructed using WGCNA v1.69 (Langfelder and Horvath, 2008; Langfelder and Horvath, 2012) using log-transformed reads in counts per million (CPM) obtained in R via the cpm function edgeR v3.35.0 (Chen et al, 2016; McCarthy et al, 2012; Robinson et al, 2010).

Genes with fewer than two read counts for four or more samples were removed from the datasets to reduce spurious correlations. The Pearson correlation metric was used to calculate expression similarity before a signed adjacency matrix was constructed with a soft-threshold power of 16 for *Arabidopsis* samples and 24 for *Lotus*. The topological overlap and topological overlap dissimilarity matrix were calculated from each species’ network adjacency matrix and used to perform average linkage hierarchical clustering with a dynamic tree cutting algorithm to generate modules. Correlation coefficients were calculated between the eigengenes of each module and relevant growth conditions (*i*.*e*., nutrient level) or gene expression to identify module-trait relationships.

Gene Ontology enrichment was performed with PANTHER (Mi et al. 2013). GO slimming to produce figure 4B used the following sequence: heirarchical clusters for GO terms in each module were generated in PANTHER and the most-specific term in each cluster was collected. The top 3 most-significant (FDR-adjusted p-value) and most-enriched (fold enrichment) terms in each module were collected and combined, then subjected to semantic similarity clustering in reviGO with a cutoff of 0.5 to produce a list of 25 representative terms (Supek et al 2011).

Differential expression analysis was performed in R using the edgeR package (Liao, Smyth, and Shi 2014; Robinson, McCarthy, and Smyth 2010). The estimateGLMCommonDisp function was used to test for differential expression among pairwise treatment groups and significance was evaluated based on the Benjamini-Hochberg FDR adjusted p-value <0.0. Cross-comparison of gene lists from differential expression or network analysis was executed with the VLOOKUP formula in Excel. Figures 3 and 4B were generated in R; all other plots and figures were created in Excel and Powerpoint.

**Figure S1:**
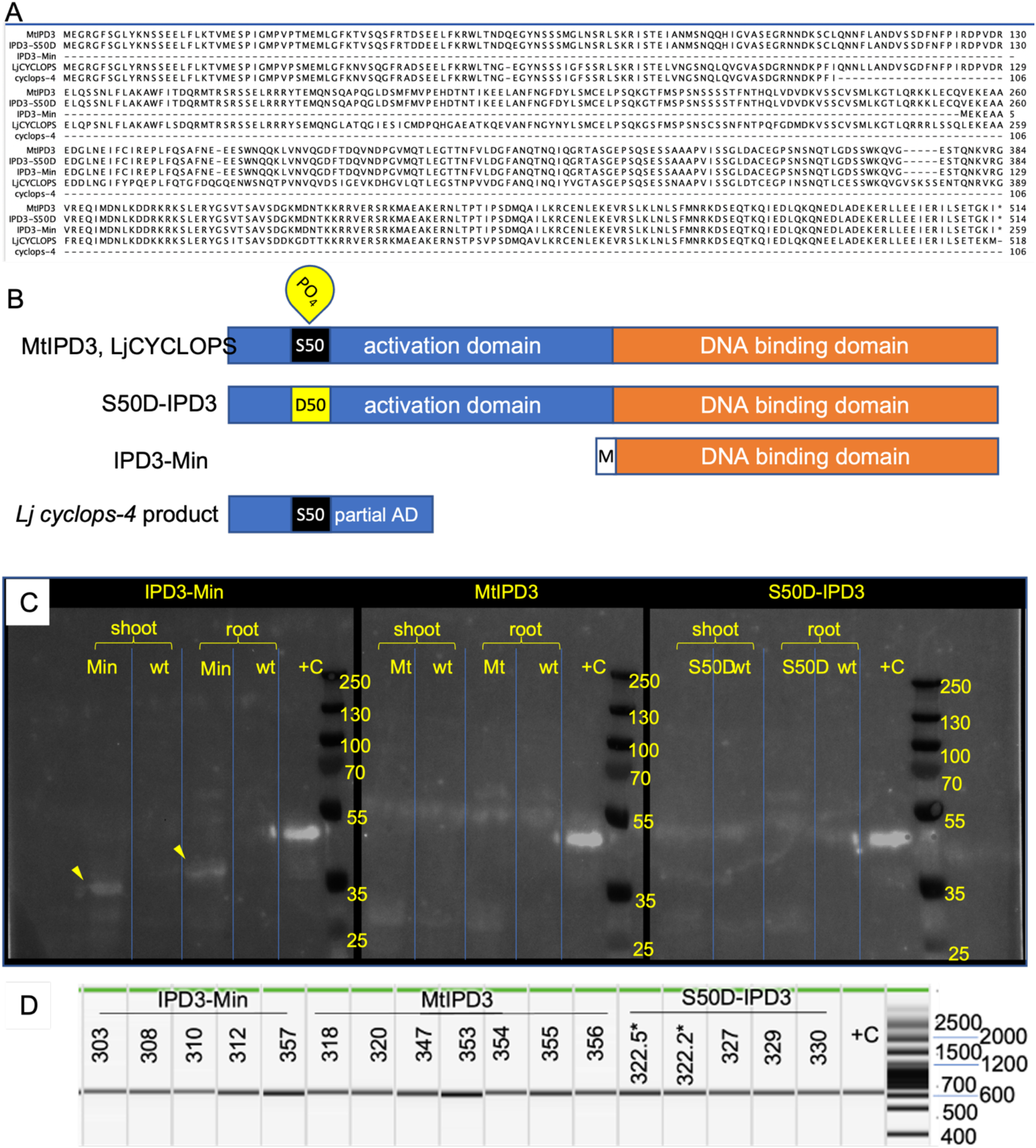
Comparison of IPD3 protein versions used in this study and validation of RNA and protein expression in transgenic plants. (A) Protein sequence alignment of *Medicago truncatula* IPD3, S50>D phosphomimic IPD3, cons=tu=vely ac=ve DNA binding domain IPD3-Min, *Lotus japonicus* IPD3/CYCLOPS, and the truncated C-terminal fragment resulting from the *cyclops-4* mutation in *Lotus*. (B) Schematic of protein domains as affected by variants in this study. (C) Western blot of 3 *IPD3* versions from transgenic *Arabidopsis* roots and shoots. The +C control sample is a synthe=c pep=de of the 349 C-terminal residues of MtIPD3 with expected size 39.98 kDa; expected size of IPD3-Min is 29.19 kDa and expected size of full-length IPD3 versions is 57.98 kDa. (D) rt-PCR to confirm expression of *IPD3* versions as RNA in leaves of T3 individuals.

**Figure S2:**
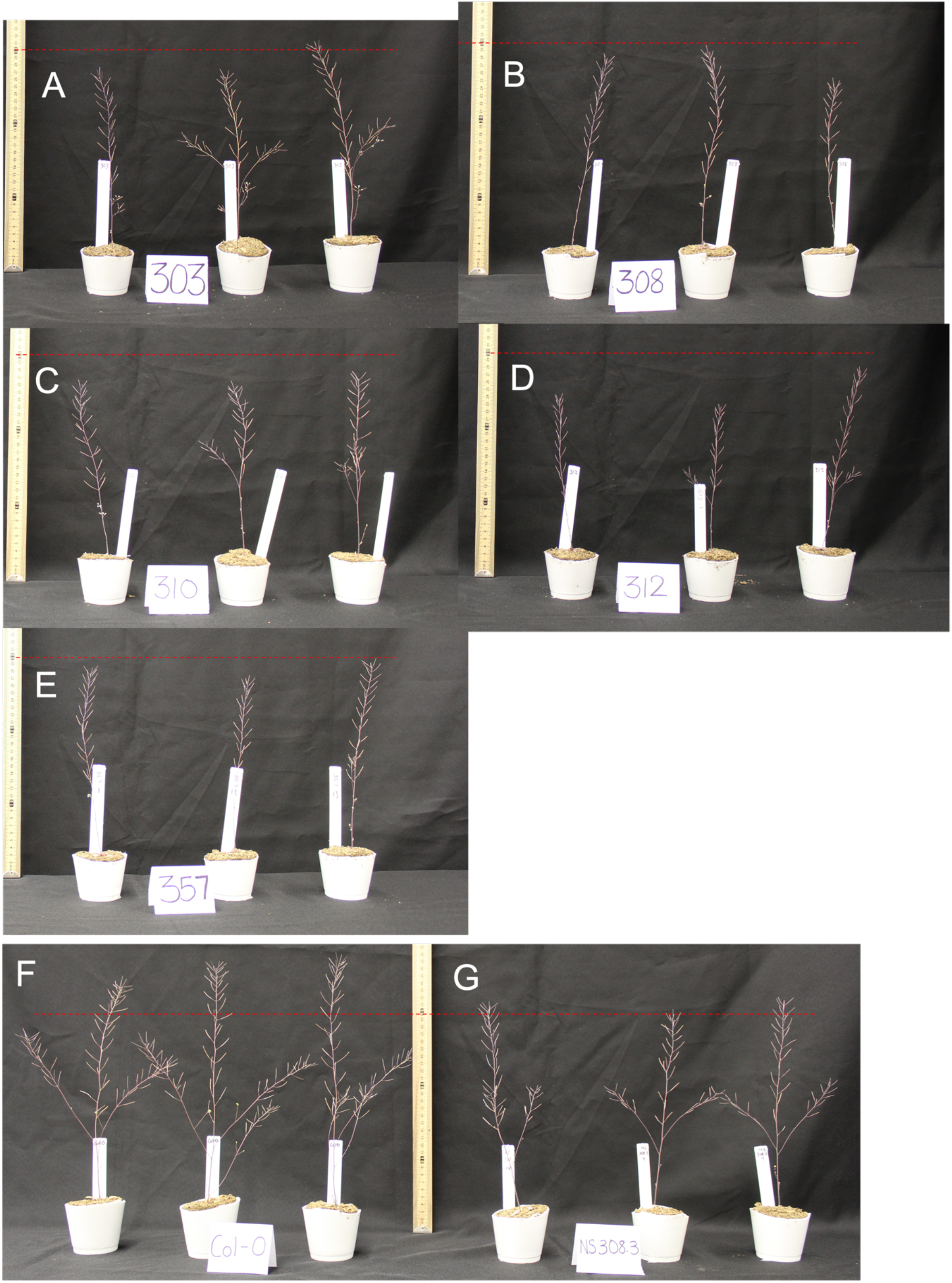
mature phenotypes of *IPD3-Min* transgenic and control *Arabidopsis* lines in T3. (A-E) Transgenic lines 303, 308, 310, 312, 357, respectively; (F) wild type Col-0; (G) null segregants of transgenic line 308. Red line in all pictures shows 30 cm from surface.

**Figure S3:**
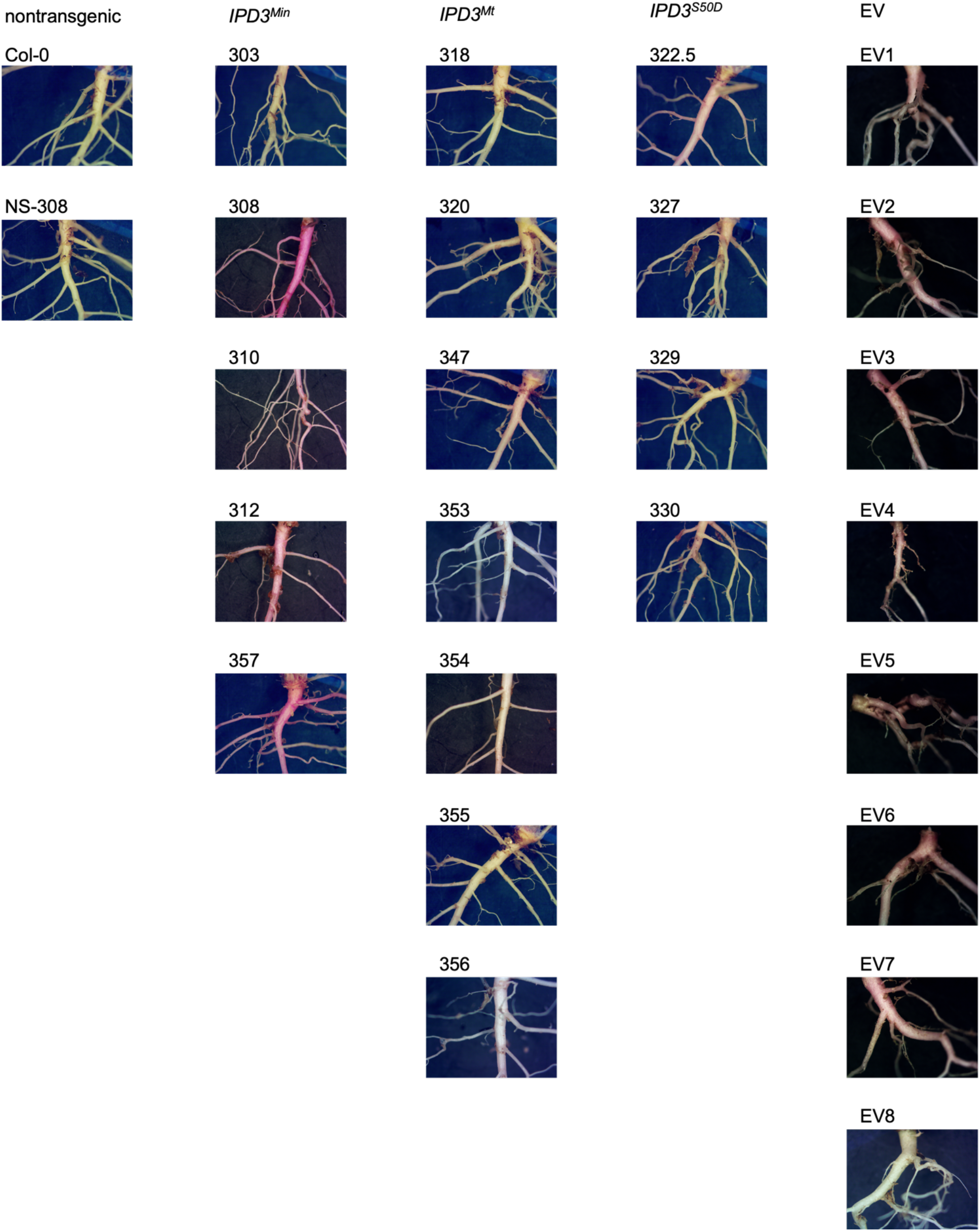
Representative root images of transgenic and control lines. In an original set of experiments, the roots of soil-grown *IPD3^Min^* transgenic plants were bright pink while wild type and null segregant controls were white, and *IPD3^Mt^* and *IPD3^S50D^* were faint pink to white. In a followup experiment, multiple independent empty vector controls containing the mCherry marker also ranged from pink to white.

**Figure S4:**
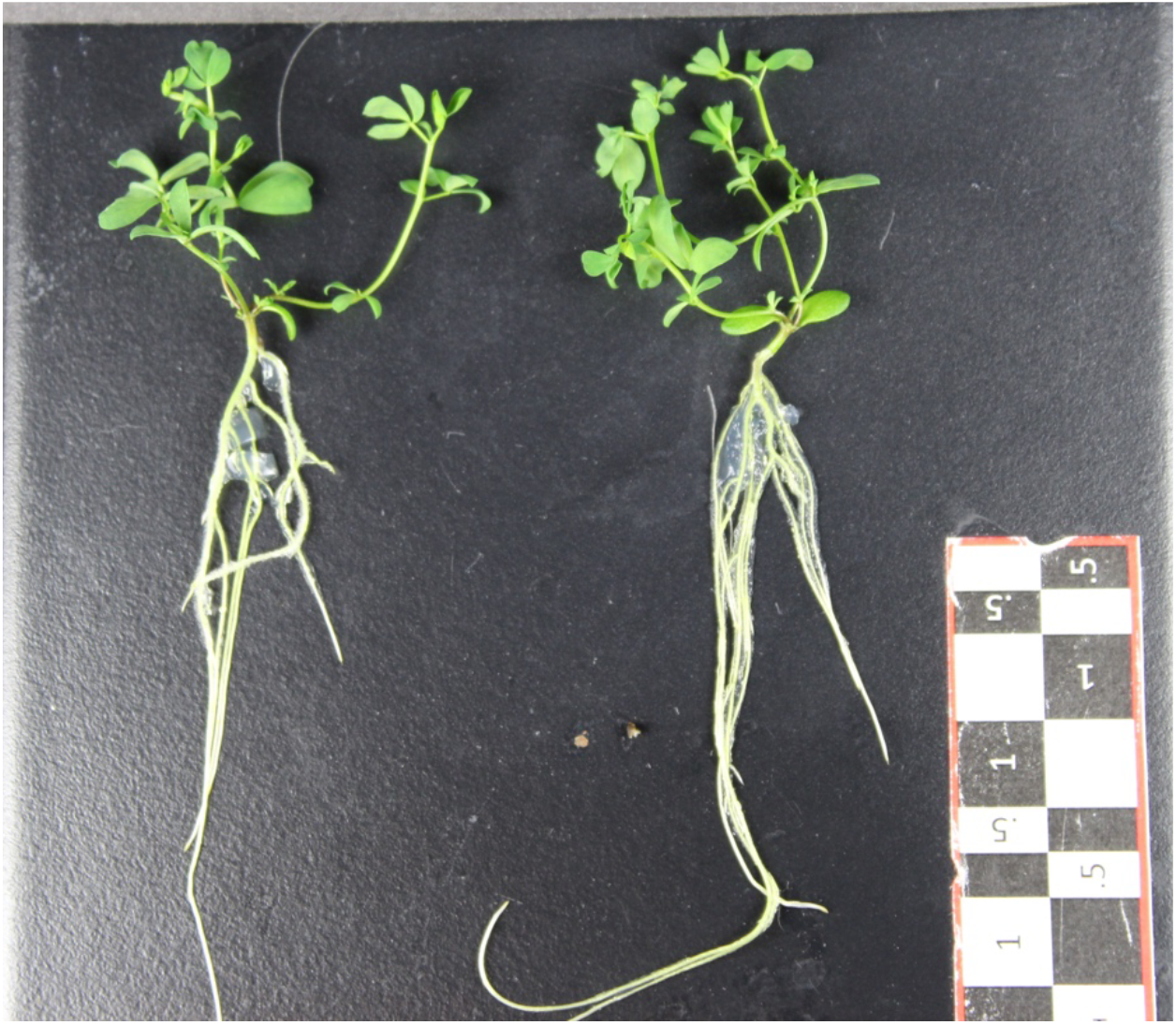
images of 5-week-old *Lotus japonicus* seedlings of Gifu wildtype and *cyclops-4* knockout mutant of *ipd3* as grown on sterile petri dishes for the transcriptome experiment. Scale bar = 2 cm (scale =les within image are denoted in cm).

**Figure S5:**
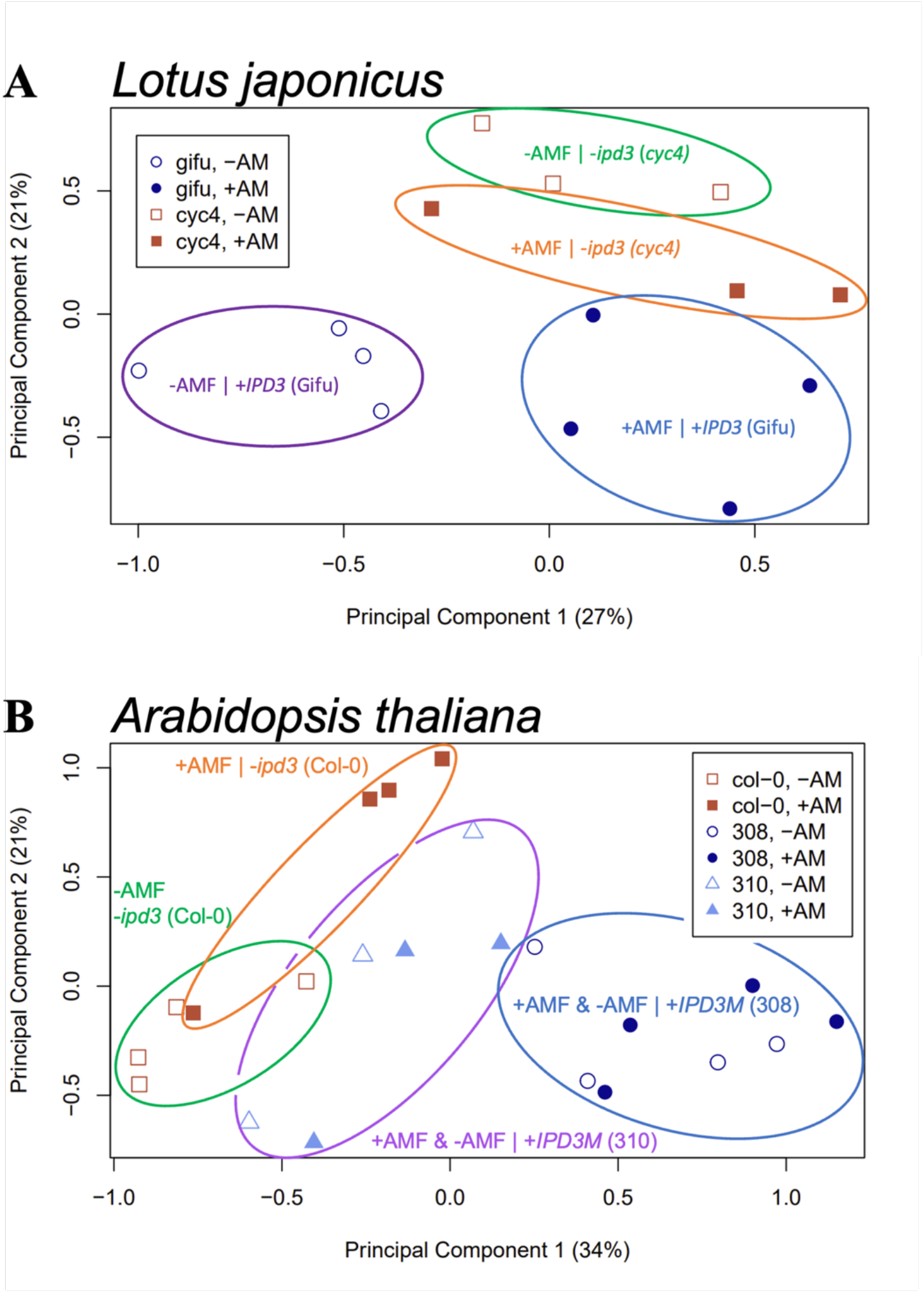
Principal component analysis shows clustering of transcriptomes by *IPD3* genotype and AMF treatment in two species. (A) *Lotus* Gifu samples cluster according to AMF treatment along PC1. Gifu and *cyclops-4* samples cluster by genotype along PC2. In contrast to Gifu, *cyclops-4* plants do not separate along PC1 according to AMF treatment; both treatment groups instead cluster near AMF-treated Gifu plants along this axis. (B) Arabidopsis samples cluster by genotype along PC1 (*IPD3M* = *IPD3^Min^* transgenic). While wild type Col-0 plants cluster by AMF treatment along PC2, there is no equivalent separation by AMF treatment within either *IPD3^Min^* line (308; 310).

**Figure S6:**
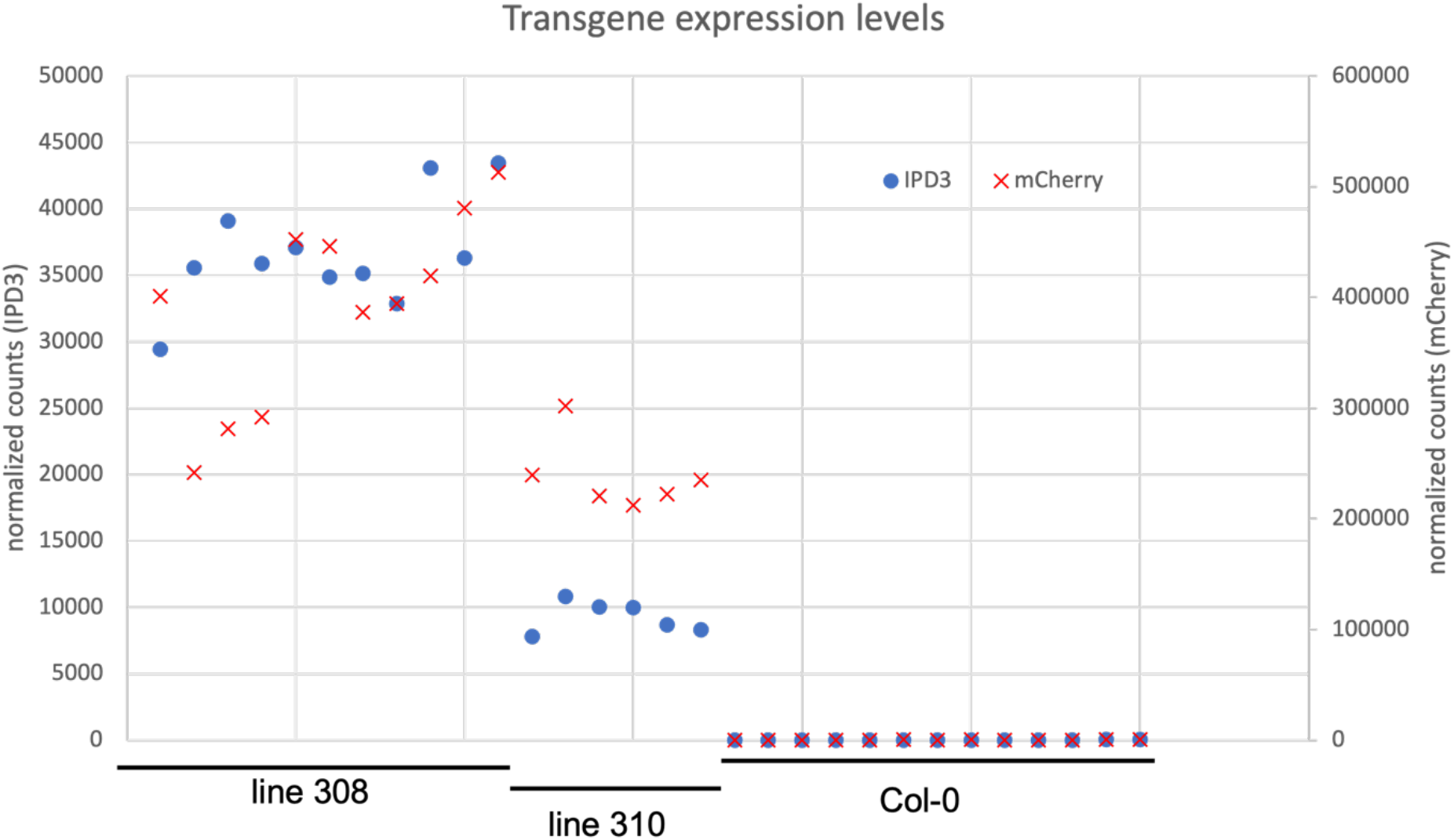
I*P*D3*-Min* and *mCherry* transcript level in *Arabidopsis* transcriptome samples. Average *IPD3-Min* transcript count in line 308 is 36,591, ∼4X higher than line 310 with average *IPD3-Min* count of 9,250.

**Figure S7:**
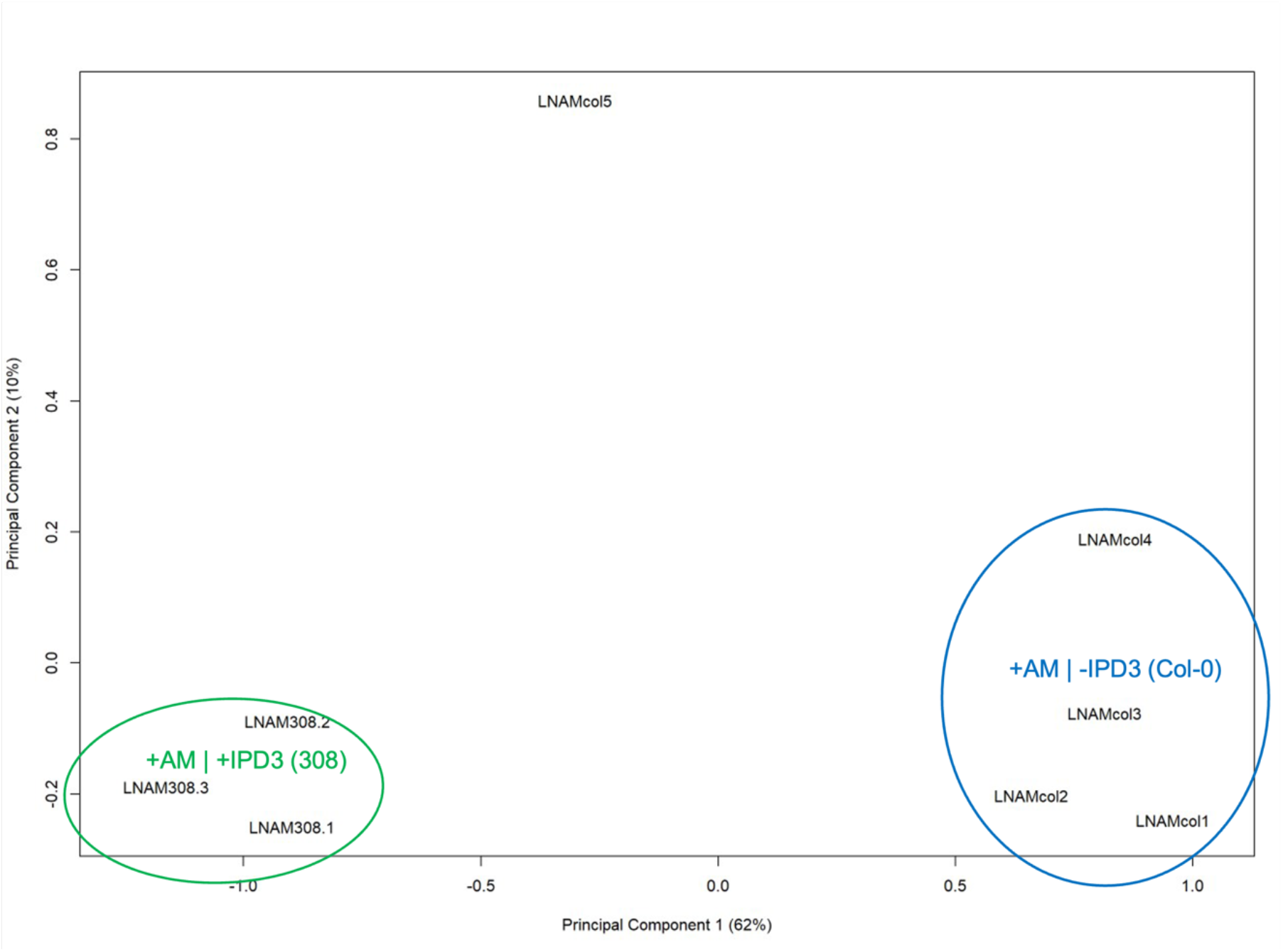
Principal Component Analysis of *Arabidopsis* transcriptomes under low-nutrient conditions with AMF treatment. Only *IPD3^Min^* transgenic line 308 was used for the low-nutrient experiment. Transcriptomes cluster by IPD3 genotype along PC1, which accounts for 62% of variation.

**Figure S8:**
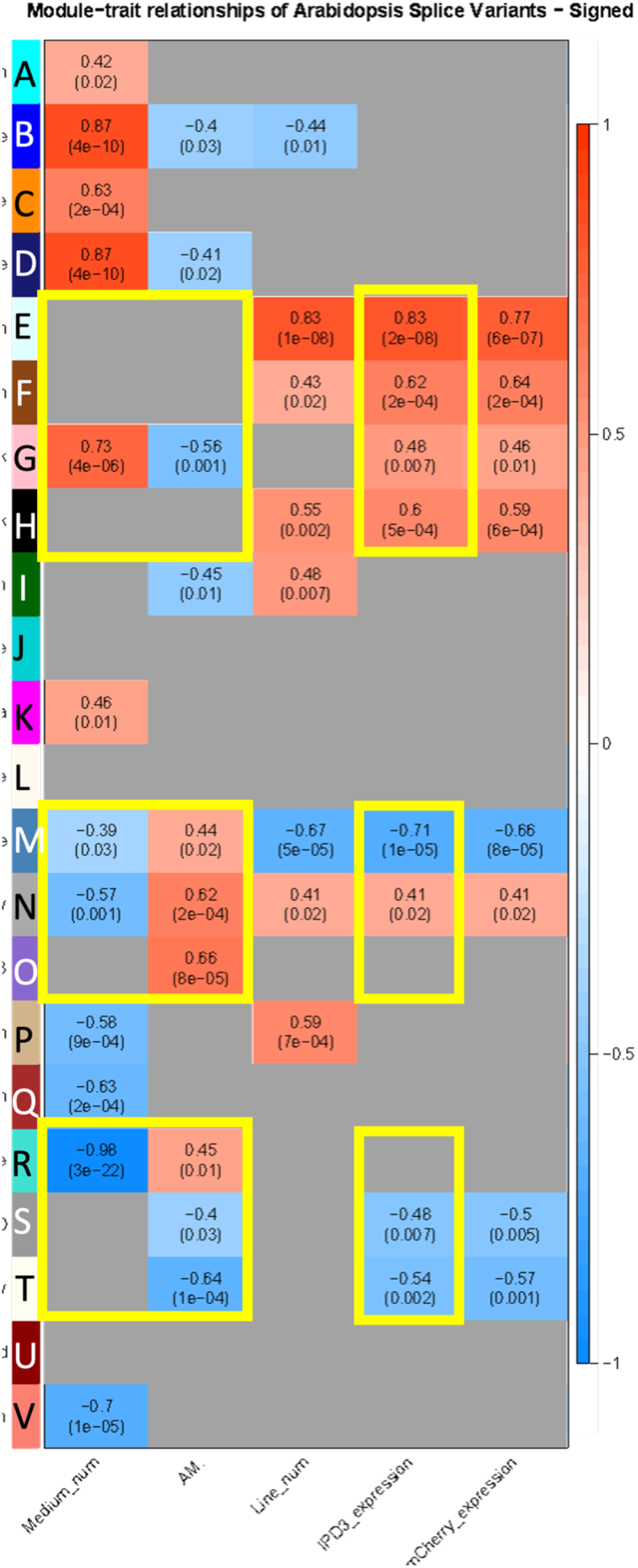
Heatmap of complete gene expression correlation network (WGCNA) with module-trait correlations. Cells included in figure 3 of the main text are highlighted in yellow. The top number in each cell is the signed eigengene correlation of that module to value of trait shown on the X axis. A value of 1 indicates perfect posi=ve correla=on, -1 would indicate perfect inverse correla=on. Boiom value shown in parentheses is the P-value of that module-trait correlation.

**Figure S9:**
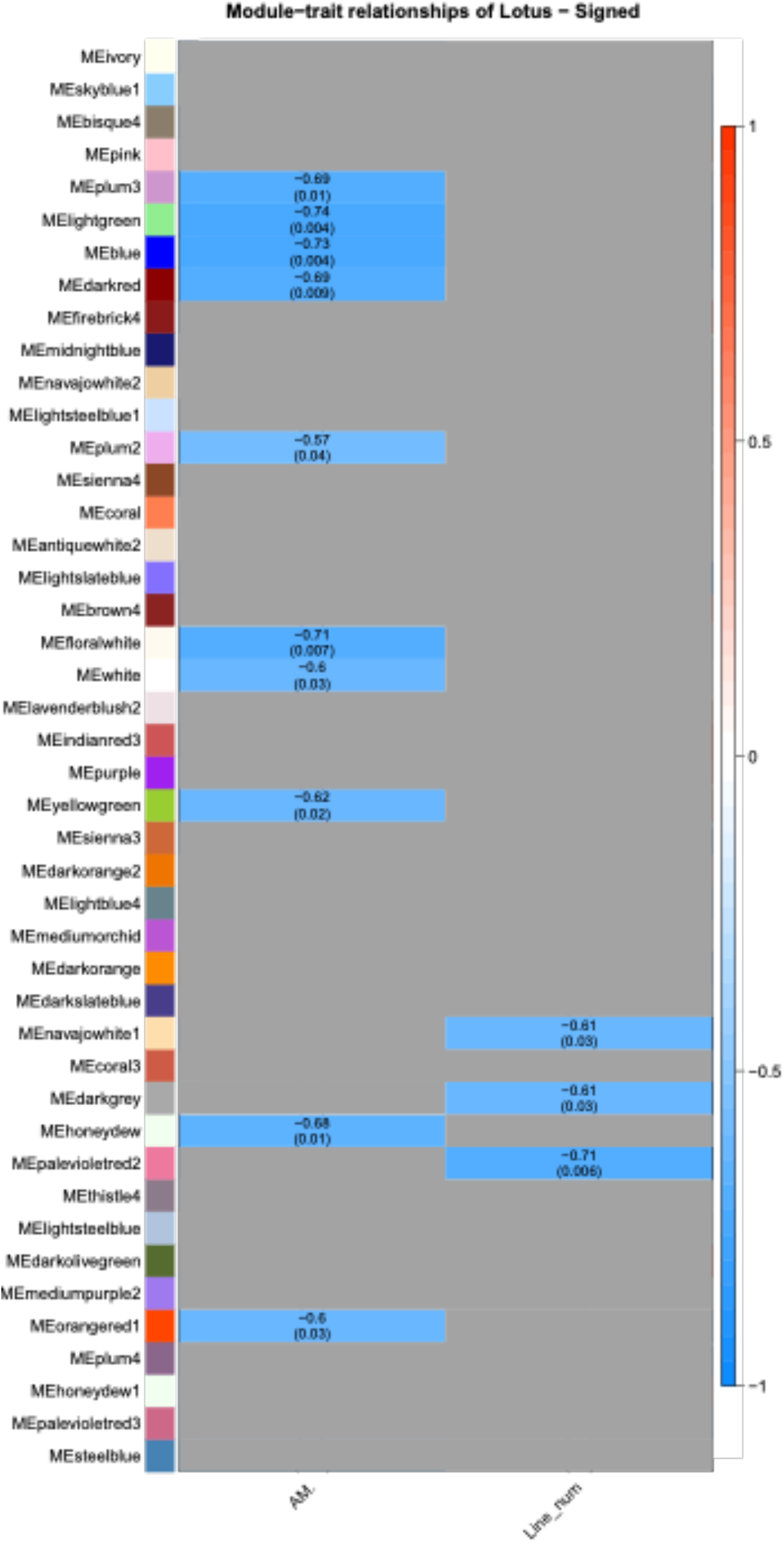
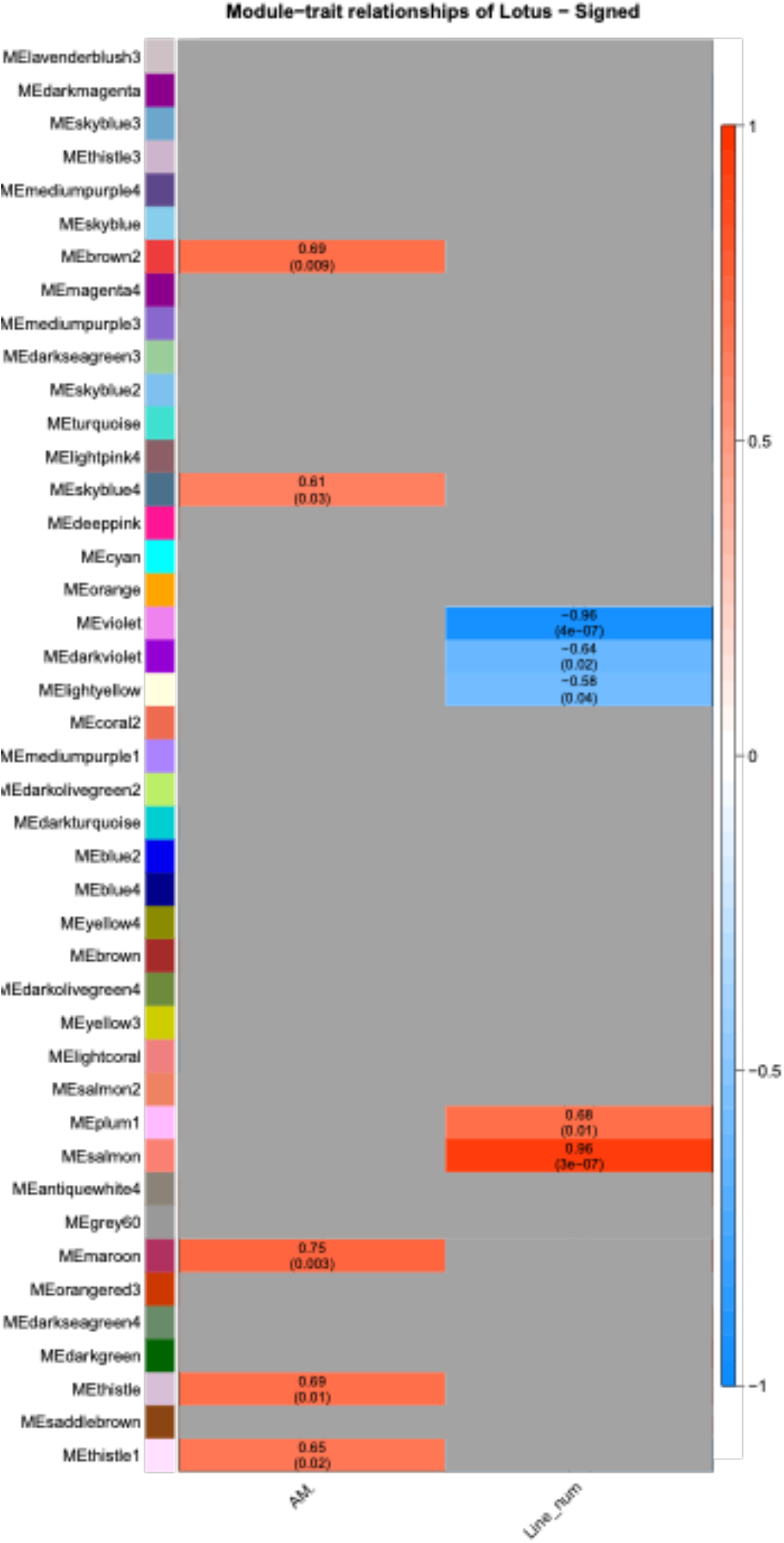
Gene co-regulatory modules resulting from WGCNA analysis of *Lotus* transcriptomes. AMF corresponds to AM fungus treatment. Geotype reflects the presence of functional *IPD3* in the wild type with a value of +1 and its absence in the *cyclops-4* mutant with a value of 0.

**Figure S10:**
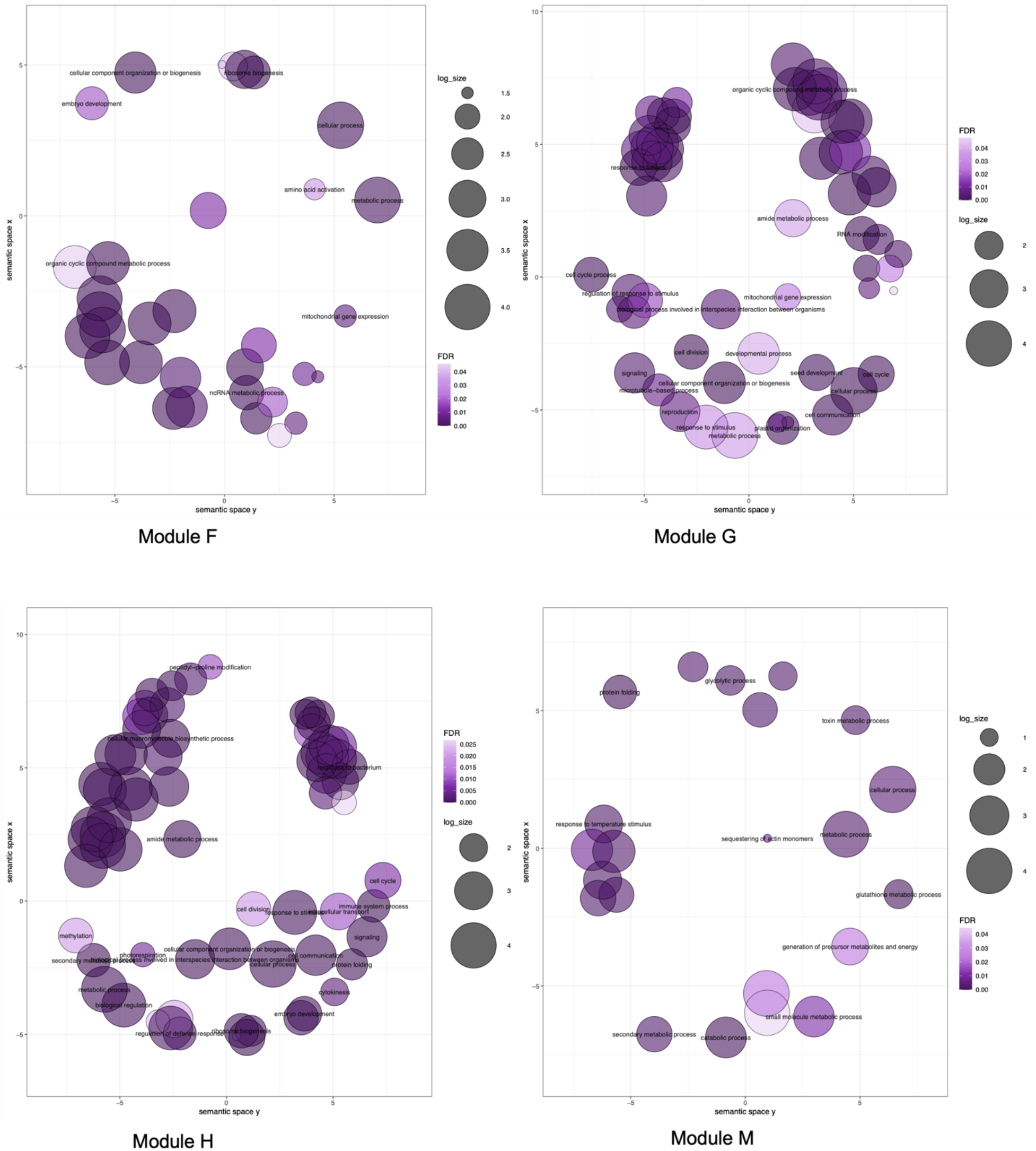
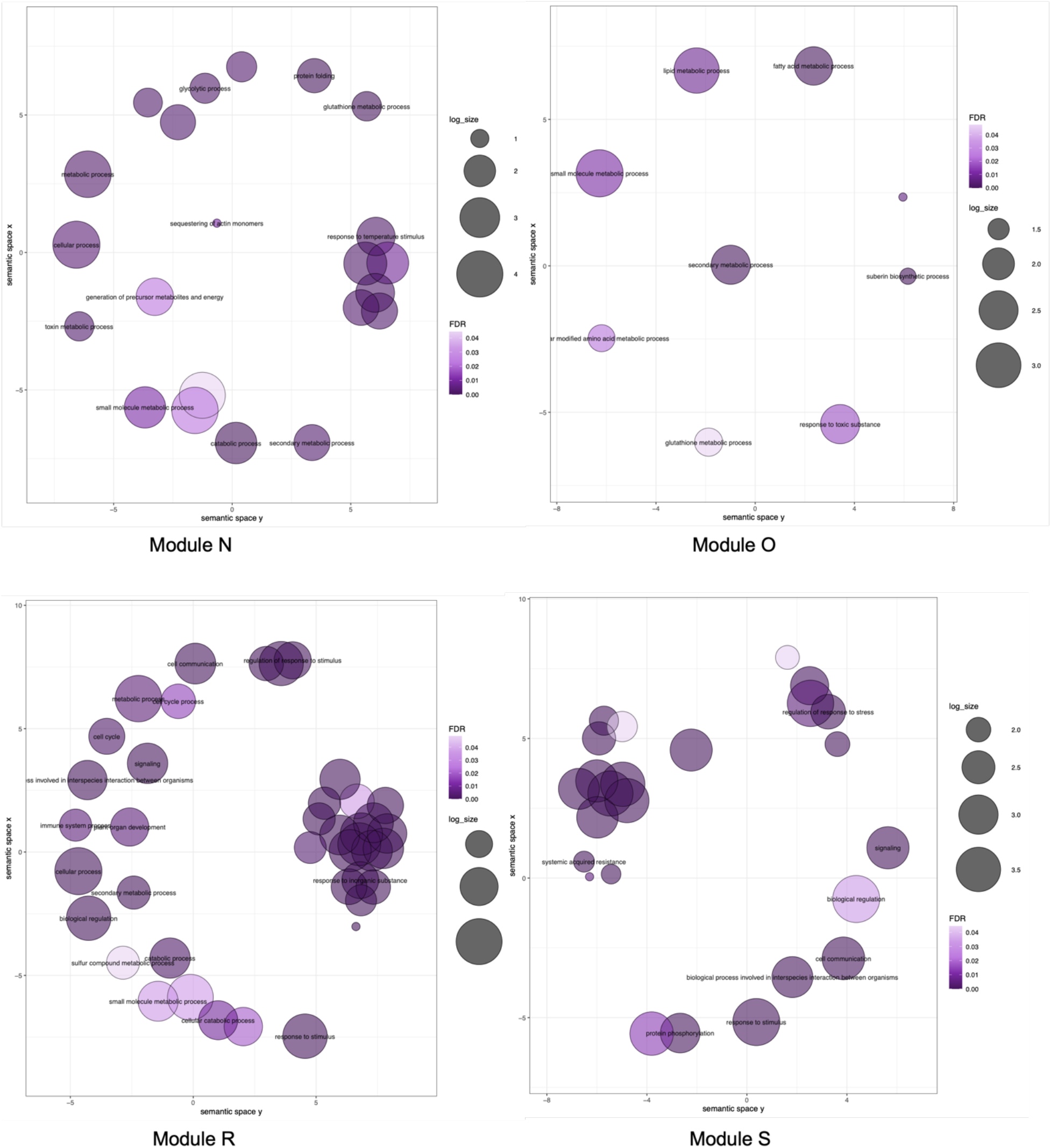
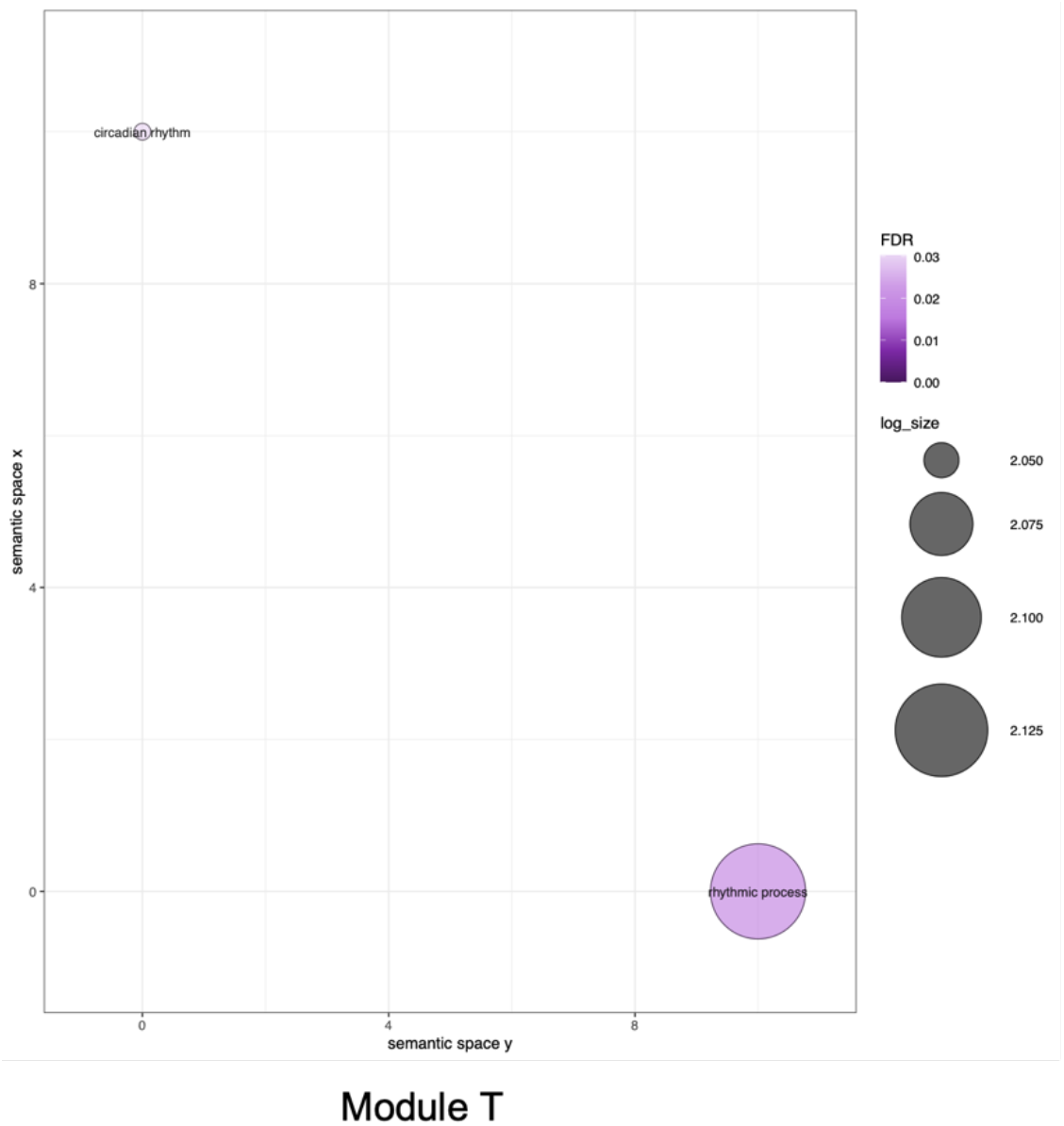
Seman=c similarity clustering of gene ontology (GO) terms of all modules shown in figure 3 of the main text (including all *Arabidopsis* modules significantly correlated to *IPD3^Min^* expression). Size of the point on the plot for each GO term corresponds to its fold enrichment in that module, and intensity of color corresponds to FDR-corrected p-value of the enrichment. Proximity of the points represents their semantic similarity as determined by ReviGO (Supek et al 2011) from relatedness and frequency within the *Arabidopsis* gene ontology reference.

## Notes

https://www.ncbi.nlm.nih.gov/geo/query/acc.cgi?acc=GSE225213

### Competing Interest Statement

The authors have declared no competing interest.

### Summary of Updates

Figures 1 and 2 and most supplemental figures did not appear correctly in the previous version due to a PDF conversion error. The PRIDE accession number for proteomics data has been added to the Data Availability section.

